# Sporadic distribution of a new archaeal genetic code with all TAG codons as pyrrolysine

**DOI:** 10.1101/2024.09.30.615893

**Authors:** Veronika Kivenson, Samantha L. Peters, Guillaume Borrel, Aleksandr Kivenson, Leah T. Roe, Noah X. Hamlish, Khaled Fadhlaoui, Alanna Schepartz, Simonetta Gribaldo, Robert L. Hettich, Jillian F. Banfield

**Affiliations:** Innovative Genomics Institute, University of California, Berkeley, CA, USA; Biosciences Division, Oak Ridge National Laboratory, Oak Ridge, TN, USA; Institut Pasteur, Université Paris Cité, Unit Evolutionary Biology of the Microbial Cell, Paris, France; Independent Researcher, New York, NY, USA; Department of Chemistry, University of California Berkeley, Berkeley, CA, USA; Department of Molecular and Cell Biology, University of California Berkeley, Berkeley, CA, USA; Université Clermont Auvergne, CNRS, Laboratoire Microorganismes : Génome et Environnement, Clermont-Ferrand, France; Université Clermont Auvergne, UMR 454 MEDIS UCA-INRAE, Clermont-Ferrand, France; Chan Zuckerberg Biohub, San Francisco, CA, USA; Arc Institute, Palo Alto, CA, USA; Earth and Planetary Science, University of California, Berkeley, CA, USA; Department of Environmental Science Policy, and Management, University of California, Berkeley, Berkeley, CA, USA; Earth and Environmental Sciences, Lawrence Berkeley National Laboratory, Berkeley, CA, USA; Biomedicine Discovery Institute, Monash University, Clayton, VIC, Australia

## Abstract

Numerous genetic codes developed during the evolution of Eukaryotes and three are known in Bacteria, yet no alternative genetic code has been established for Archaea. Some bacterial and archaeal proteins include selenocysteine or pyrrolysine, the 21^st^ and 22^nd^ amino acids, but no evidence establishes the adoption of a genetic code in which a stop codon universally encodes either amino acid. Here, we used proteomics to confirm the prediction that certain Archaea consistently incorporate pyrrolysine at TAG codons, supporting a new archaeal genetic code which we designate Genetic Code 34. This genetic code has 62 sense codons encoding 21 amino acids, and only two stop codons. In contrast with monophyletic genetic code distributions in bacteria, Code 34 occurs sporadically. This, combined with evidence for lateral gene transfer of the code change machinery and anticipated barriers to code reversal, suggests Code 34 arose independently in multiple lineages. TAG codon distribution patterns in Code 34 genomes imply a wide range in time since code switch. We identified many new enzymes containing Pyl residues, raising questions about potential roles of this amino acid in protein structure and function. We used five new PylRS/tRNA^Pyl^ pairs from Code 34 archaea to introduce new-to-nature pyrrolysine analogs into proteins in *E. coli*, demonstrating their utility for genetic code expansion.

**Graphical abstract:** 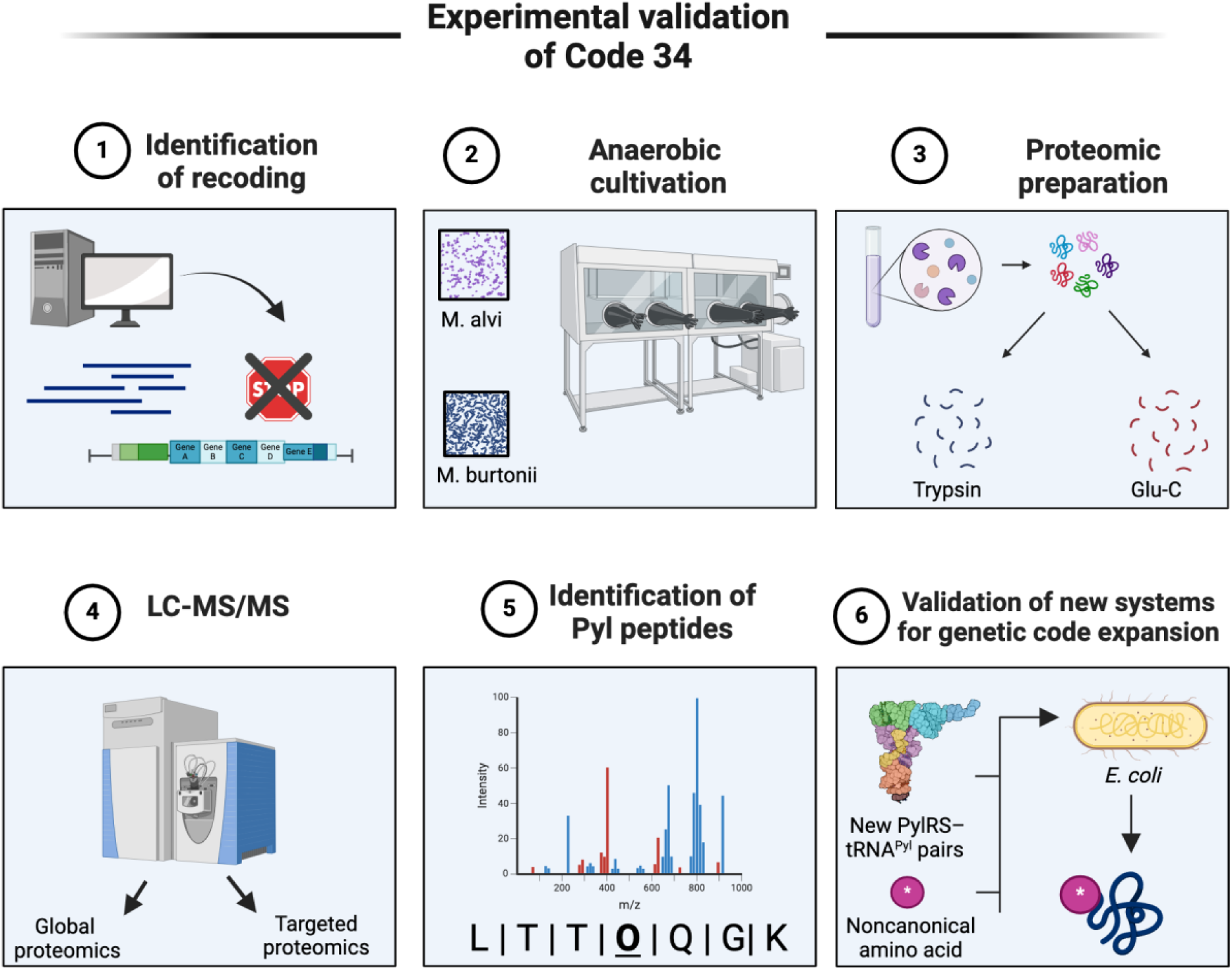

## Main

Although viewed historically as a frozen accident in time^1^ the genetic code is neither universal nor immutable. Subsequent to its initial discovery, analysis of genetic material from diverse lineages revealed substantial code variation, and even instances where different codes are used within a single organism (e.g., by mitochondria vs. the nuclear genome). More recently, new genetic codes in which either TGA or TAG stop codons incorporate a standard amino acid were identified in bacteria of two phyla and in certain bacteriophages^2–7^. Interestingly, these stop codons can also be selectively interpreted as either selenocysteine (the 21^st^ amino acid, at TGA) or pyrrolysine (the 22^nd^ amino acid, at TAG)^8,9^. Pyrrolysine (Pyl) is a derivative of lysine that features a pyrroline ring^9,10^. Its biosynthesis requires the Pyl B,C,D genes while its incorporation into proteins requires a pyrrolysyl-tRNA synthetase (PylRS) and a Pyl tRNA carrying a CUA anticodon^9,11–14^. We refer to the Pyl B,C,D genes and PylRS/tRNA^Pyl^ pair as the Pyl machinery.

PylRS/tRNA^Pyl^ pairs are widely used tools for genetic code expansion (GCE), in which non- canonical ⍺-amino acids or their analogs are incorporated into ribosomal products to modify protein function^15^. PylRS/tRNA^Pyl^ pairs are particularly useful for GCE because they act on ⍺- amino acid substrates with diverse side chains and are orthogonal in many bacterial and mammalian systems, that is, they show limited reactivity with canonical ⍺-amino acids and neither interfere nor integrate with prokaryotic or eukaryotic aminoacyl-tRNA pairs.

PylRS/tRNA^Pyl^ pairs also remain active towards substrates with non-canonical ⍺-substituents, notably ⍺-hydroxy acids^16^ and ꞵ^2^-hydroxy acids^17^, resulting in proteins with site-specific edits within the backbone itself. The introduction of ⍺-hydroxy acids is especially useful, as it facilitates more widespread backbone edits^18^, including the introduction of polyketide-like elements^19^. Yet the yields of proteins engineered using known PylRS/tRNA^Pyl^ pairs vary substantially as a function of protein identity, incorporation site, and substrate type. As a result, there remains a need for new PylRS/tRNA^Pyl^ pairs that support the synthesis of novel and diverse hetero-polymers for biotechnology applications.

In organisms that naturally encode the Pyl machinery, understanding of the mechanism that signals Pyl incorporation is incomplete and the extent to which Pyl occurs in proteins remains unclear. It has been suggested that a PYLIS, a structure analogous to the SECIS required to selectively signal selenocysteine incorporation, enables a dual meaning of the TAG codon^20,21^. However, later analyses revealed that the PYLIS is absent in a subset of genes that encode Pyl proteins^22,23^. In bacteria, Pyl has been reported only in one L-serine dehydratase and a few methylamine methyltransferases^9,11,24,25^ and otherwise, TAG is read as a stop codon. Pyl has been predicted to occur in proteins of 11 major groups of archaea^9,26–36^. All archaea without Pyl machinery use TAG as a stop codon and those with the Pyl machinery are believed to use TAG as a stop codon, except in the very few enzymes in which Pyl is known to occur. Additionally,

Pyl has been hypothesized to occur within proteins when internal TAG codons were observed during *in silico* analysis^26,37,38^. However, incorporation has been experimentally confirmed only in mono-, di-, tri- methylamine methyltransferases, tRNA guanylyltransferase, and PylB^10,22,39,40^.

The archaeon *Methanosarcina acetivorans* has five additional proteins presumed to contain Pyl because their detected molecular masses are consistent with TAG readthrough^22^. The authors suggest that TAG may not be selectively reassigned in *M. acetivorans*^22^, but genome-wide incorporation of Pyl at this codon was not established.

We asked if some archaea broadly incorporate Pyl into proteins using an alternative genetic code, and whether these genomes would yield new biotechnologically relevant PylRS/tRNA^Pyl^ pairs. Our analysis revealed that organisms within two major groups of Archaea contain multiple genes with internal in-frame TAG codons. Using global and targeted proteomics, we analyzed two such Archaea, a human gut and a marine isolate, and confirmed in both cases the genome-wide incorporation of Pyl at TAG codons, including within multiple new enzyme classes. On this basis we propose the existence of a previously unrecognized genetic code, currently confined to archaea, and designate it as Genetic Code 34. We confirmed that five new PylRS/tRNA^Pyl^ pairs predicted by our analysis efficiently introduce Pyl analogs into proteins in *E. coli*, some with unique specificities. This work confirms that Genetic Code 34 organisms can add unique “parts” to the genetic code expansion toolbox and facilitate the synthesis of new-to-nature heteropolymers. We find that the distribution of Genetic Code 34 among archaea is not monophyletic, which is consistent with multiple independent code transitions and hints that codon reassignment to Pyl could be facile. We infer features indicative of recoding that occurred recently vs. in the distant past and propose a series of steps that may lead to genetic code transition.

### Unusual phylogenetic distribution of Pyl

To investigate the phylogenetic distribution of archaea with Pyl machinery (Pyl-encoding archaea; Pyl archaea), we augmented a collection of genomes from all families known to incorporate Pyl, together with new taxa that we determined carry the Pyl machinery (**Table S1, S2**). The majority of Pyl archaea belong to the Halobacteriota (Methanosarcinia, Archaeoglobi and Methanonatronarchaia) and Thermoplasmatota (Methanomassiliicocci) phyla, with few others within the phyla of Asgard, Thermoproteota, and Hydrothermarchaeota (**Fig. 1, Table S3**). Notably, organisms that encode Pyl machinery have a discontinuous phylogenetic distribution (**Fig. 1**), sometimes even at the genus level. For example, the Pyl pathway is encoded and actively utilized by a single member of the Archaeoglobus (*Ca.* Methanoglobus hypatiae LCB24)^28^. The proteins from this genome share 98.6% average amino acid identity (AAI) with Archaeoglobi LCB024-003^28^, yet LCB024-003 does not encode Pyl. In a similar way, Pyl is limited to a single family within the Asgard and Hydrothermarchaeota phyla, and in the Ca. Methanomethylicus genus, only two of five archaea encode Pyl.

**Fig. 1.**
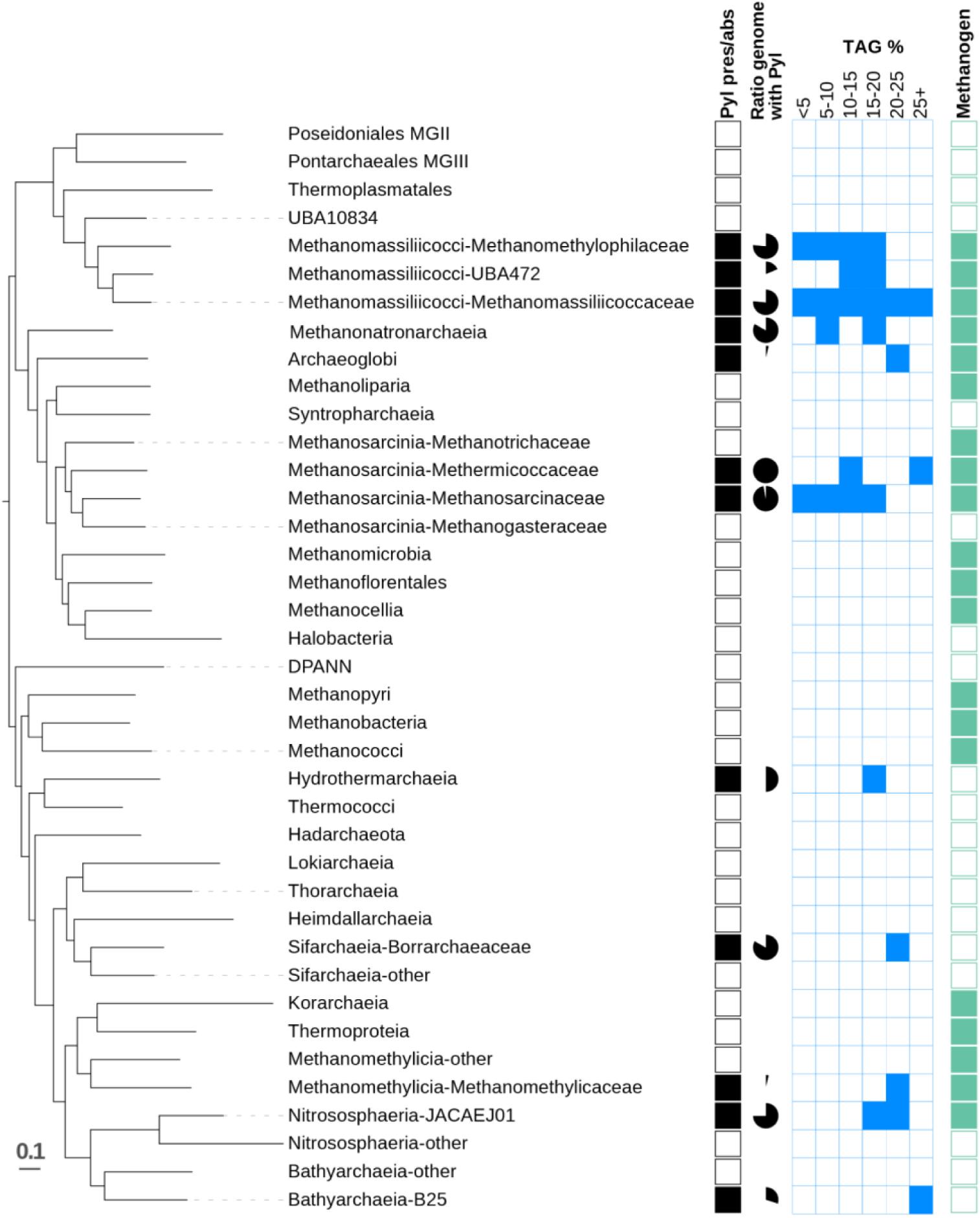
Distribution of the Pyl machinery in the Archaea is paraphyletic, with the Pyl machinery variably present among representatives of the same family. The range of TAG frequency of Pyl genomes (blue boxes) in each lineage illustrates that low TAG frequency is not a hallmark of Pyl archaea, in contrast to previous reports.

Given the intermixing of closely related archaea that do and do not encode the Pyl machinery, we compared the phylogenies of the Pyl machinery (**Fig. 2**) to that of the organisms (**Fig. 1**). Consistent with prior suggestions^41,42^, discordance between the tree structures provides evidence for lateral transfer of the Pyl genes. For example, MSBL1 archaea (Hadarchaeota) likely gained their Pyl machinery through horizontal gene transfer from Methanonatronarchaeia, and “*Ca.* Methanoglobus” from a Thermoproteota.

**Fig. 2.**
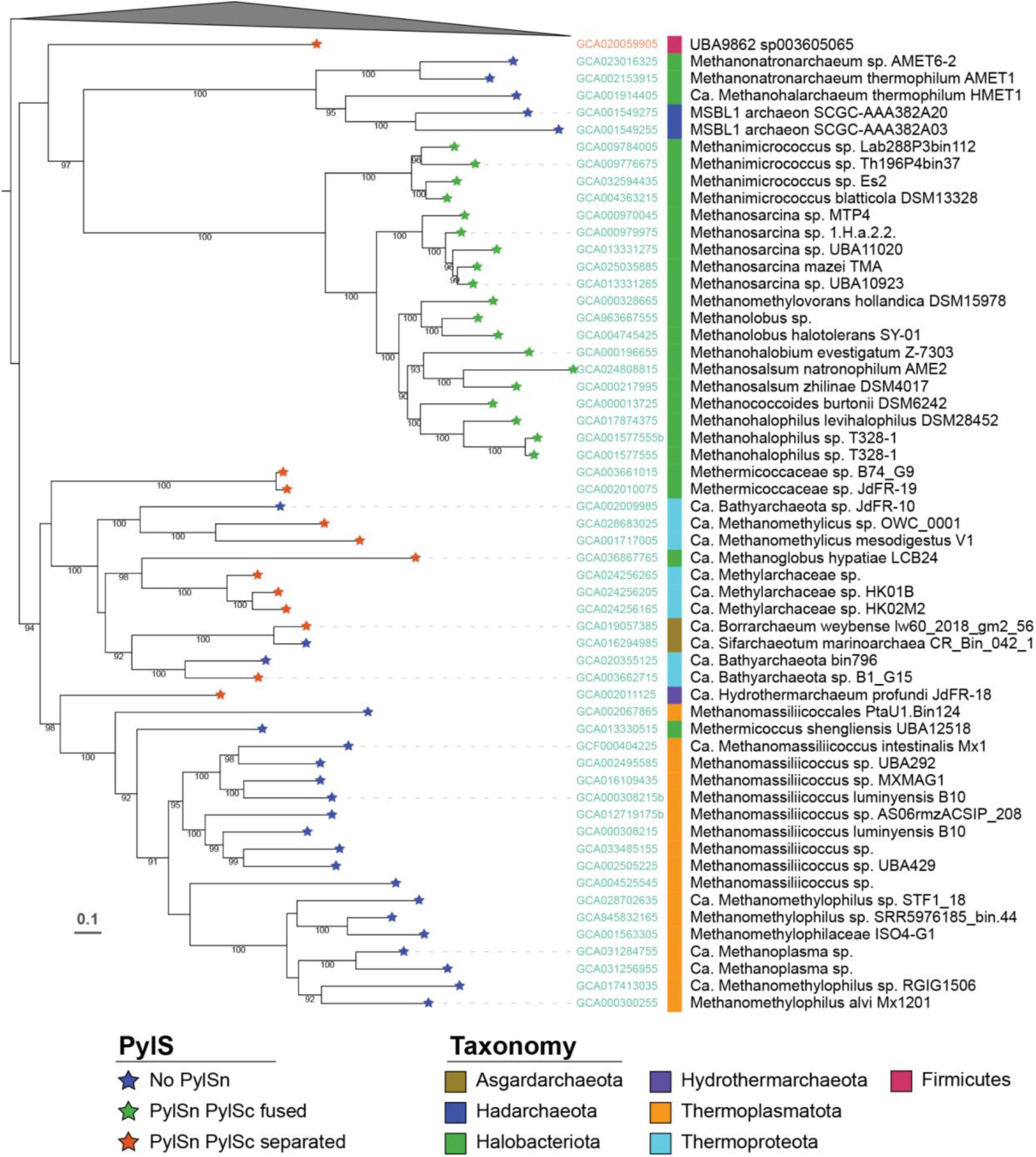
Maximum likelihood phylogeny of the PylBCDS enzymes. Stars on the branch indicate the status of PylSn (missing, fused with PylSc or separated from PylSn). 1,241 positions, ultrafast bootstrap values >90% are displayed.

## TAG frequency varies among Pyl archaea

Given that organisms almost always interpret TAG as a stop codon^43^, we reasoned that there may be detrimental consequences associated with the acquisition of a Pyl machinery, such as the erroneous incorporation of Pyl and readthrough of the normal gene end. It is also possible that TAG codons possess a dual meaning in organisms that carry a Pyl machinery, however there exist no validated mechanisms for deciphering a dual-message TAG codon. Initial studies suggested that the presence of an RNA stem loop immediately downstream of the TAG codon (a Pyl insertion sequence, or PYLIS) is required to interpret a TAG codon as Pyl and not as stop^20,21^. However, the presence of a PYLIS was later shown to have no effect on levels of Pyl incorporation in *E. col^i^*^44^ and no PYLIS or PYLIS-like structures could be detected in a subset of known Pyl-containing methylamine methyltransferase genes in archaea^22,23^. We examined sequences of genes encoding the Pyl-containing tRNA guanylyltransferase and PylB, and no PYLIS was detected (**Table S4**).

One strategy that would avoid or minimize the need to decipher a dual-message TAG codon is simply to eliminate its use as a stop codon. In fact, it has long been thought that low TAG frequency (≤ 5%) represents a hallmark of archaea that encode a Pyl machinery^23,24^, yet a few outliers have been reported^26,29,33^. We calculated the percentage of genes with a TAG codon in the genomes of Pyl archaea, and find that the TAG frequencies are always high (>15%) among Pyl archaea that are intermixed phylogenetically with genomes that lack the Pyl machinery (**Fig. 1, Table S3**). Thus, we conclude that low TAG frequency is not a core feature across Pyl archaea. The discrepancy from expected TAG content in many Pyl archaea highlights the need to determine the distribution of TAG codons and investigate how they are used.

## TAG is recoded genome-wide as a sense codon

To determine how TAG is used by Pyl archaea, we first predicted the genes in each organism using Genetic Code 11, which is standard for bacteria and archaea. We then re-predicted all TAG codons as sense codons and classified the resulting genes into four categories (**Fig. 3, Supplementary Fig. 1**). Category 1 genes contain one or more TAG codons in which the sequences before and after the TAG codon possess >30% sequence identity to a known protein (see *Methods*). In addition, readthrough of the TAG codon in Category 1 genes extends the protein sequence by >15 amino acids (aa). Based on this analysis, we predict that genes in Category 1 interpret the TAG as an internal sense codon. Category 2 genes were characterized by short extensions of ≤15 aa, and had no overlap with sense codon(s) of an adjacent gene.

**Fig. 3.**
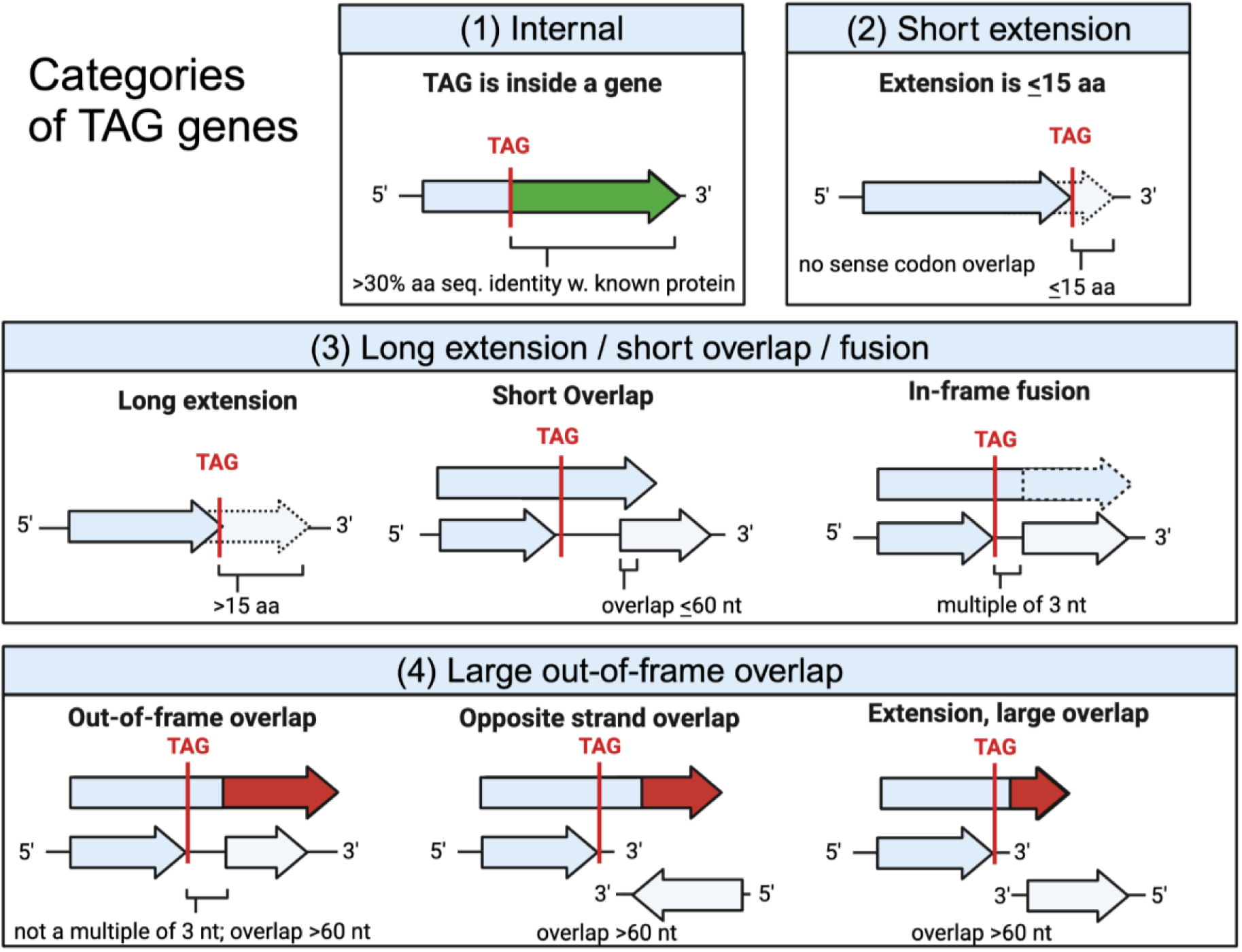
TAG-containing genes in Pyl-encoding archaea are sorted into one of four categories based on the result of readthrough of the TAG codon. Category 1 (internal) genes contain an internal TAG codon, and encode proteins for which the amino acid sequence before and after the TAG codon has >30% sequence identity with sequences from the UniProt databases. Category 2 includes short extensions, when sequence extension after TAG results in a ≤15 aa extension and no sense codon overlap. Category 3 includes genes with extension past TAG resulting in an in-frame fusion, a short overlap with an adjacent gene, or a long extension. Large-out-of frame overlaps (>60 nt) are assigned to Category 4.

Genes in Category 3 were characterized by an extension of >15 aa after TAG readthrough (sequence extension lacks similarity with known genes) and/or a short (<60 bp) gene overlap and/or in-frame gene fusion. For Category 2 and 3, it is not possible to predict whether TAG is a stop or sense codon. For Category 4, readthrough led to a large (>60 nt) out-of-frame overlap. In this case, there is no support for TAG reassignment, as large gene overlaps (>60 bp) are rare in prokaryotes^45^. The prevalence of Category 1 and absence or extreme rarity of Category 4 cases in a genome may be an indicator of genome-wide TAG reassignment.

Notably, Pyl archaea from eight genera (Methanomethylophilus, Methanogranum, Methanoplasma, Methanimicrococcus, Methanosarcina, Methanohalobium, Methanohalophilus, and Methanococcoides) have numerous Category 1 TAG genes (**Fig. 4**, **Table S5**). In fact, in a subset of the Thermoplasmatota and Euryarchaeota, most TAG codons are found within Category 1 genes and there are very few, or no, Category 4 genes. This pattern is consistent with reliance only on TGA and TAA as stop codons. We infer that in these genomes, TAG is completely recoded as a sense codon (Pyl). Thus, we sought experimental evidence to confirm that these archaea interpret the TAG codon as Pyl.

**Fig. 4.**
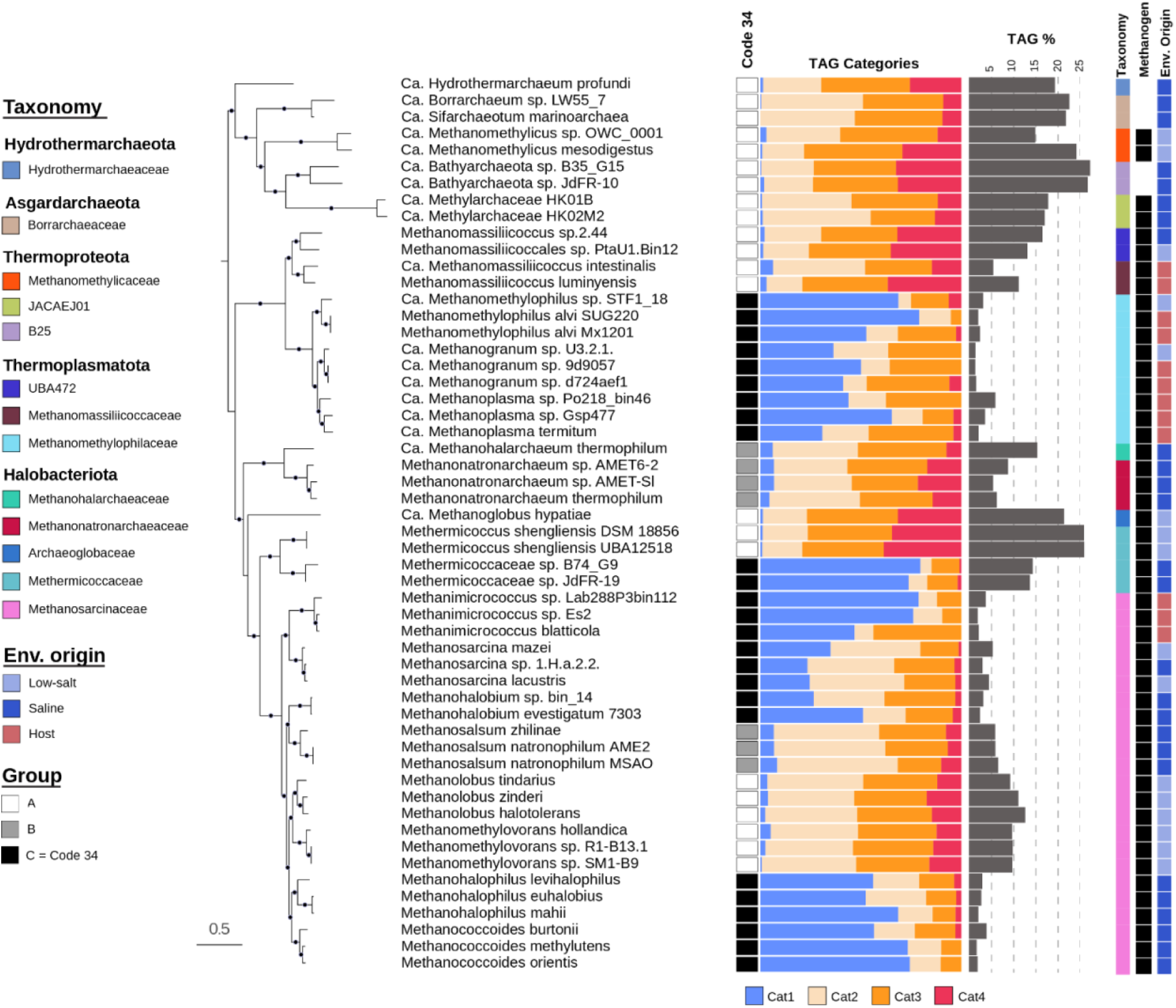
Phylogenetic tree showing evidence for the sporadic distribution of Genetic Code 34 in archaea. Genomes inferred to be fully recoded (black boxes, Group C) have a large fraction of internal TAG codons (Category 1) and very few cases where TAG readthrough leads to large out-of-frame gene overlaps (Category 4). In Group A genomes, TAG content is high (>15%), while Group B genomes typically have 5 - 15% TAG content and internal TAG in genes in addition to the methylamine methyltransferases. Within Group C, the TAG content is variable, probably in part due to time since code transition. Tree constructed using 40 universally conserved proteins. 9816 positions, ultrafast bootstrap values >90% are displayed.

## Proteomic validation of Genetic Code 34

To test for recoding of TAG, we chose two isolates from different lineages and environments with genomes that are predicted to contain a large fraction of Category 1 genes and used mass spectrometry to identify proteolytic peptides and their subsequent fragmentation whose exact masses confirmed the presence of a Pyl residue. *Methanococcoides burtonii* is a psychrophilic Halobacteriota from an Antarctic lake^46^ and *Methanomethylophilus alvi* (formerly *M. alvus*) is a Thermoplasmatota from the human gut^31,47^. To increase the probability of detection of the specific peptides of interest, we used both trypsin and endoproteinase Glu-C, which cleave between different amino acids and thus increased the number of peptides suitable for analysis. Pyl-containing tryptic or GluC-fragments will be observed if and only if two criteria are met: the Pyl-containing protein is sufficiently abundant, and the Pyl-containing fragment(s) possesses the appropriate mass and charge to be detected. Although comprehensive detection of all peptides is not possible due to variable abundance, searching specifically for peptides of interest based on their predicted masses (targeted peptide detection) greatly improved our ability to identify Pyl-containing peptides (see *Methods*). Peptides with the mass-to-charge ratio (m/z) expected for a Pyl-containing tryptic or GluC-product were manually validated by analysis of the MS/MS spectra.

For both *M. burtonii* and *M. alvi*, the proteomic data confirm genome-wide recoding of TAG as a sense codon (**Table S6, S7**). The detection of ≥2 unique peptide matches in the global tryptic or Glu-C datasets was used to confirm protein expression (**Table S6, S7**). Notably, for all proteins that were expressed and predicted to contain Pyl, and for which a proteolytic peptide (>6aa and <50aa) spanning the region predicted to contain Pyl was detected, the presence of Pyl was confirmed. For *M*. *burtonii,* peptides containing Pyl were detected in 27 of 39 expressed Pyl proteins. Similarly, for *M. alvi*, Pyl was detected in 12 of the 15 expressed Pyl proteins.

Expressed proteins in which Pyl was not detected had low sequence coverage and / or had a Pyl-containing proteolytic peptide that was too short or too long for detection. As an additional confirmation of Pyl incorporation at TAG codons, additional de-novo-assisted database searches were conducted to enable identification of any of the standard amino acids at the TAG codon. We did not detect any peptides with another amino acid in the TAG position in the *de novo*-assisted database searches. Pyl was detected in 24 additional proteins that had low levels of expression (<2 unique peptides matches in the global datasets). In total, Pyl was detected in 55 proteins not previously shown to contain Pyl (**Table S8**).

Examples of new proteins found to contain Pyl in *M. burtonii* include 12,18- didecarboxysiroheme deacetylase, thiouridine synthase, DNA helicase, histidine tRNA synthetase, isocitrate dehydrogenase, epimerase, glycosyltransferase, DNA polymerase, and multiple hypothetical proteins (**Fig. 5A, Supplementary Fig. 2A, Table S6**). New Pyl proteins in *M. alvi* include transacetylase-like protein, a methyltransferase, kinase, acetyltransferase, hypothetical and DUF-containing proteins, and a AAA family associated protein (**Fig. 5B, Table S7**). Pyl was also identified in trimethylamine methyltransferase (**Supplementary Fig. 2B**), as previously reported^10,40^. Thus, we conclude that *M. burtonii* and *M. alvi* archaea have adopted a previously undefined genetic code, which we propose as Genetic Code 34, with 62 sense codons encoding 21 amino acids, and only two stop codons. It is probable that the other Pyl-encoding archaea for which we detected evidence for numerous internal TAG codons (Category 1) also use Genetic Code 34.

**Fig. 5.**
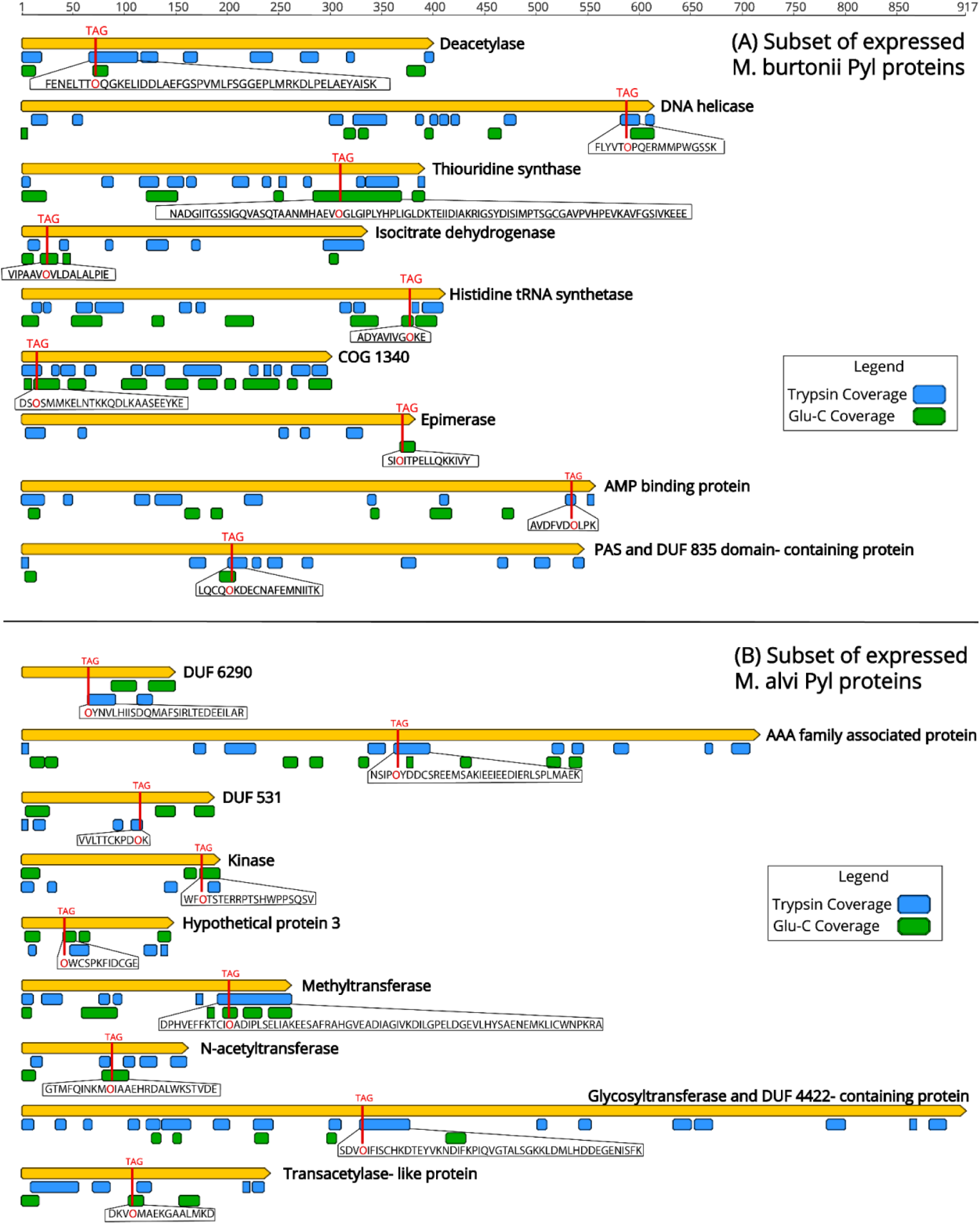
Proteomic coverage confirms Pyl residues and extensions past the TAG codons for proteins expressed by **a,** *M. burtonii* and **b,** *M. alvi*. Coverage based on enzymatic cleavage using Trypsin is shown in blue and Glu-C is shown in green. The TAG location is shown with a vertical red line and Pyrrolysine is denoted by the one-letter designation, “O”.

## Over a thousand proteins are predicted to contain Pyl

After establishing that select archaea use TAG as a dedicated sense codon, we predicted 1903 unique Pyl-containing proteins in genomes inferred to use Code 34 (**Table S9**). This finding increases the number of predicted Pyl proteins by two orders of magnitude. Of the 1903 proteins, Codetta^48^ predicted that 360 contain TAG within the gene based on alignment of profile hidden Markov models (HMMs) of conserved proteins (**Table S10**). The other cases involve N-terminal or C-terminal extensions beyond the TAG codon that do not align with known protein sequences (HMMs), or fusions with subsequent genes in the same reading frame. The largest number of predicted Pyl proteins of a single organism, 191 in total, occurs in JDFR-19 (Methermicoccaceae; **Table S11**). This archaeon is from crustal fluids collected from oceanic crust near the Juan de Fuca Ridge^49^, and is a recoding outlier with a 13% TAG frequency.

Pyl occurs in numerous protein families, but the only Pyl-containing proteins that archaea have in common are the methylamine methyltransferases. The proposed function of Pyl in these proteins is to activate and orient the methylamine for methyl transfer to a cognate corrinoid protein, via the reactivity of the electrophilic imine bond^10,40^. Outside of the methyltransferases, nearly all proteins newly found to incorporate Pyl have a limited phylogenetic range, such that Pyl is in a protein within a single genome or genus, as a singleton. These may arise due to neutral mutations, as described for Pyl in tRNA-His guanylyltransferase^39^. The position of Pyl is maintained in a GAF domain-containing / PAS family protein in three genera of the Methanosarcinaceae family (**Supplementary Fig. 3**). In methylcobalamin:CoM methyltransferase (MtaA), Pyl is in a different position in the amino acid sequence in organisms belonging to two genera (positions 90 and 289 in Methanosarcina and Methanohalophilus respectively) but both positions are predicted to be near the opening of the binding cavity in the folded protein (**Fig. 6**). It remains to be established what (if any) role Pyl plays in the enzymology of these new Pyl-containing proteins.

**Fig. 6.**
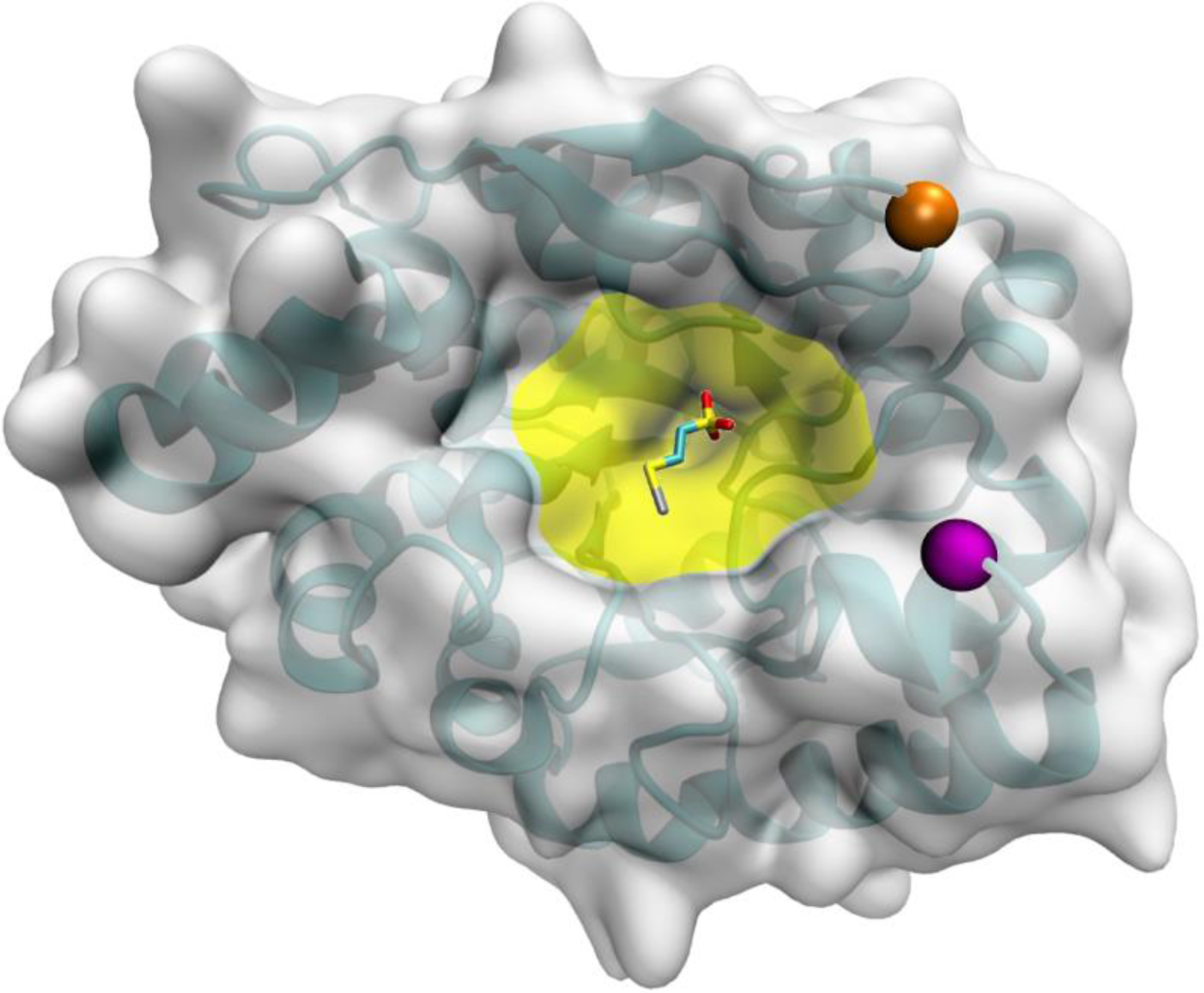
Pyl residues without a conserved sequence position in related genes may still play similar roles due to their spatial proximity in the folded proteins. The positions of the Pyl residue in homologs of methylcobalamin:CoM methyltransferase (MtaA) from Methanosarcina sp. (orange) and Methanohalophilus sp. (magenta) are marked on the structure of MtaA (PDB 4AY7). Both positions are near the opening of the binding cavity (yellow, shown in complex with Zn^2+^ and coenzyme M). The actual positions may be even closer together due to potential differences in the structures of the two homologs.

Pyl occurs in many proteins involved in energy metabolism and DNA processing, including citrate synthase, isocitrate dehydrogenase, aminoacyl-tRNA synthetases, DNA polymerases, and helicases. Additionally, transposases and CRISPR Cas proteins contain Pyl. Pyl was previously predicted in transposases and the Cas1 and Cas3 proteins^26,42,50^. Here, we observed Pyl in Cas1 via targeted proteomic analysis (**Table S6**), and identified 12 new transposase families with Pyl, with some occurring in many copies per genome. One example is *Methanosarcina mazei* strain TMA, which has 49 copies of the IS66 family transposase with Pyl (**Table S10**). Pyl is also in Cas2, Cas4, Cas8e, Cas10, and there is a region with multiple recoded proteins in a Cas1 casposon (**Table S12**). Casposons are a superfamily of mobile elements that encode Cas1, family B DNA polymerase, and conserved, uncharacterized protein domains^51^. Qualitatively, this incidence is not notably different to that predicted if TAG codons reflect random mutation, so enrichment due to selection for beneficial effect in Cas proteins is not indicated. However, as TAG codons remain very rare, lack of evidence for selection may simply reflect insufficient time since code transition.

Archaeal TnpB and IscB proteins both are predicted to contain Pyl (**Table S12**). TnpB and IscB are RNA-guided nucleases that are the likely ancestors of Cas12 and Cas9, respectively ^52,53^. The location of the Pyl residue is not conserved in IscB homologs, but in each of the five homologs where the Pyl residue is in the homologous region, it is close to the heteroduplex of the guide RNA and target DNA, in either the bridge helix or the HNH endonuclease domain (**Fig. 7**). The Pyl-containing nucleases identified in this study represent a diverse and overlooked set of potential gene-editing proteins. In studies of IscB and TnpB nucleases, the Pyl-containing variants are always either mis-annotated as fragments, hypothetical proteins, or missed altogether (**Supplementary Fig. 4**), and the Pyl-containing casposon was assumed to be inactivated due to the internal TAG codons. The findings of the current study motivate experimental work to test the functions of these and other newly defined Pyl-containing proteins.

**Fig. 7.**
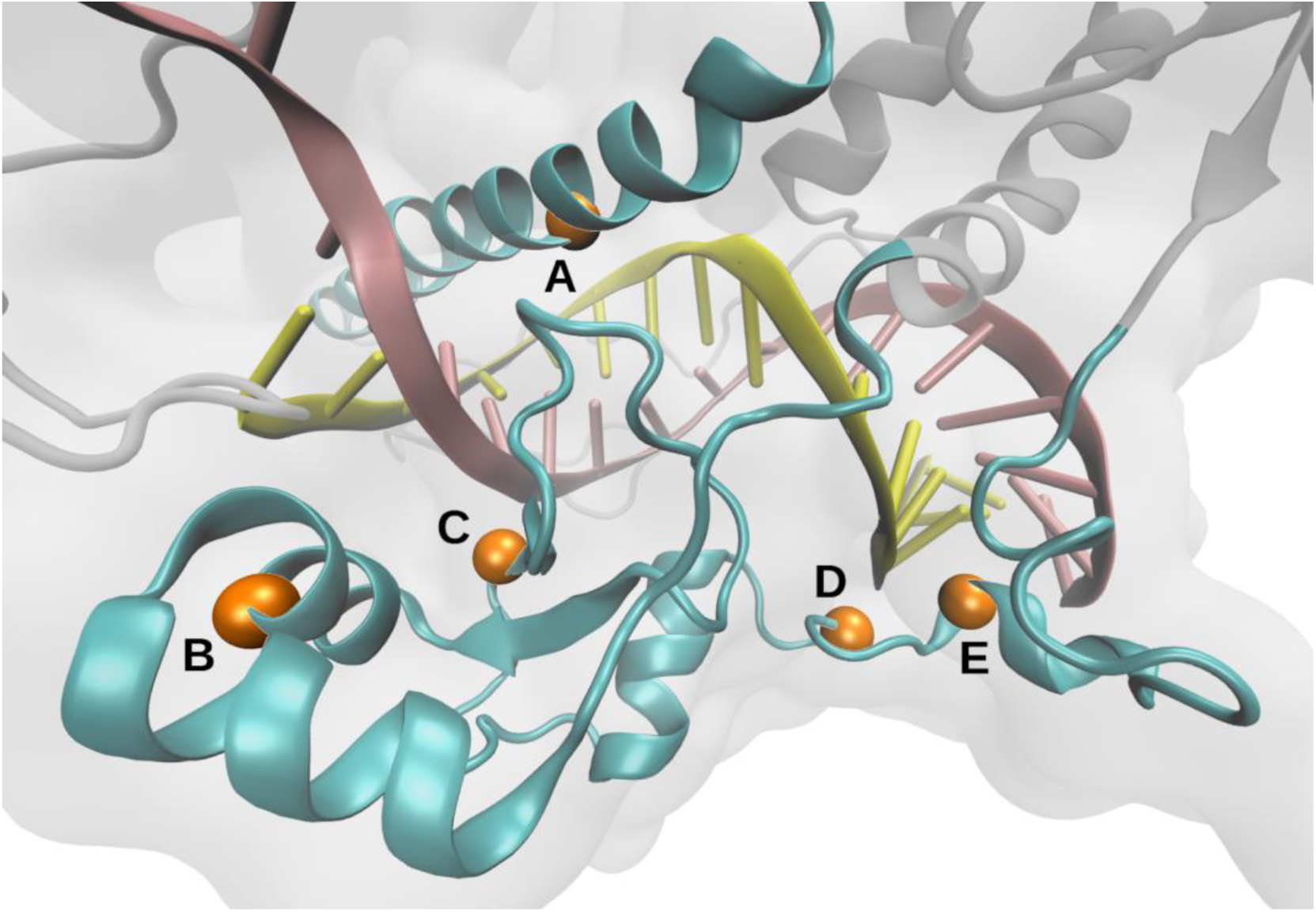
Pyl residues may play a role in the binding of IscB homologs to genetic material. The positions of the Pyl residue in each of five putative homologs of IscB from different organisms (**a**, **d,** Methanosarcina lacustris sp., **b,** Methanonatronarchaeum sp. Amet6_2, **c,** Candidatus Methanohalarchaeum thermophilum, **e,** Methanohalobium evestigatum) are marked in orange on the bridge helix **a,** and HNH endonuclease domain **b,c,d,e,** of IscB (PDB 7UTN). Although these positions are not conserved, they are all located near the paired guide RNA (yellow) and target DNA (pink) strands.

## PylRS/tRNA^Pyl^ pairs from Genetic Code 34 organisms support genetic code expansion (GCE) in *E. coli*

The high incidence of TAG codon(s) encoding Pyl within genes of some Genetic Code 34 archaeal genomes suggests a robust system for Pyl incorporation that may be leveraged for the site-specific incorporation of unnatural monomers into ribosomal products. Indeed the PylRS/tRNA^pyl^ pairs from *M. mazei*, *M. barkeri*, and *M. alvi* are already widely used for GCE in bacterial, yeast, and mammalian cells^52^. Here, we tested whether PylRS/tRNA^Pyl^ pairs from newly identified Genetic Code 34 archaea support the incorporation of two widely used analogs of Pyl (**1**), the ⍺-amino acid Boc-Lys (**2**) and the ⍺-hydroxy acid HO-Boc-Lys (**3**) (**Fig. 8A**), into a model protein, super folder green fluorescent protein (sfGFP)^53^, containing an in-frame TAG codon located at position 3 (sfGFP-3TAG). At least one representative sequence from each recoded genus was evaluated. All in all, 8 new PylRS/tRNA^Pyl^ pairs were tested for their ability to support genetic code expansion in *E. coli*, using the PylRS/tRNA^Pyl^ pair from *M. alvi* as a benchmark for current GCE efficiency levels. Initial screenings evaluated whether organisms supplemented with Boc-Lys (**2**) or HO-Boc-Lys (**3**) developed higher levels of sfGFP fluorescence (F_528_) as a function of cell growth (OD_600_) over 20 hours (h) (**Fig. 8B, Supplementary Fig. 5**). When BL21(DE3) *E. coli* were supplemented with Boc-Lys (**2**), the PylRS/tRNA^Pyl^ pairs from *Methanomethylophilus alvi* and *Methanohalophilus levihalophilus* developed substantial levels of sfGFP compared to non-supplemented growths, while the PylRS/tRNA^Pyl^ pair from *Methanococcocoides LMO-1* supported moderate sfGFP expression.

**Fig. 8.**
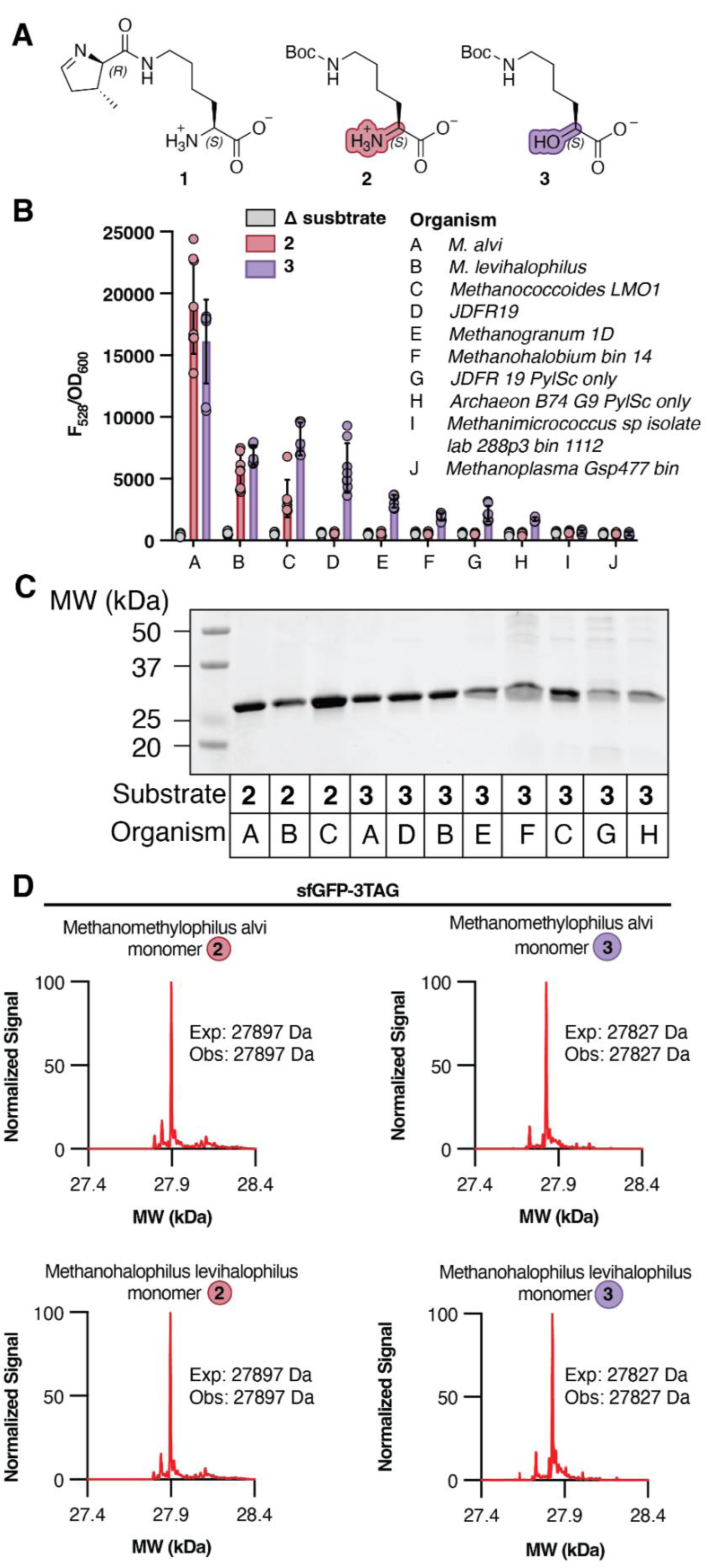
PylRS variants from organisms that utilize Genetic Code 34 support the incorporation of non-canonical ⍺-amino and ⍺-hydroxy acids into the model protein sfGFP. **a,** Chemical structures of three established PylRS substrates: pyrrolysine (Pyl, **1**) the natural substrate for PylRS; Boc-Lysine **2** (BocK), a synthetic α-amino acid substrate, and HO-Boc-Lysine **3** (BocK-HO), an α-hydroxy acid substrate. **b,** Shown is the F_528_ signal normalized to OD_600_ for bacterial cultures of BL21(DE3) cells doubly transformed with a pMega plasmid harboring the PylRS/tRNA^Pyl^ from candidate organisms A-J and a pET22A reporter plasmid containing sfGFP with an in-frame amber codon at position 3. Cells were grown to OD_600_ = ∼0.6, supplemented with 1 mM of the indicated PylRS substrate, and sfGFP expression was induced with 1 mM IPTG. F_528_/OD_600_ signal was acquired for over a 20 h time frame and the final signal at T = 20 h is shown here. We note the presence of separately encoded PylSc and PylSn domains of the PylRS protein in the JDFR19 genus. As prior studies have found PylRS enzymes naturally lacking PylSn to be active, we also evaluated 2 JDFR19 PylRS homologs with only their PylSc domain. **c,** SDS-PAGE of purified sfGFP-3TAG constructs in the presence of either 1 mM **2** or **3** and the corresponding PylRS/tRNA^Pyl^ pair. **d,** Intact MS of sfGFP-3TAG expressed in the presence of a control PylRS/tRNA^Pyl^ pair (from *M. alvi*) or in the presence of the best performing candidate PylRS/tRNA^Pyl^ pair (*M. levihalophilus*) as determined by expression tests in panel B. sfGFP was expressed in the presence of 1 mM **2** or **3**.

SDS-PAGE analysis of sfGFP-3TAG proteins expressed in BL21(DE3) *E. coli* supplemented with either 1 mM of Boc-Lys (**2**) or HO-Boc-Lys (**3**) indicates that the major protein product has a molecular mass of approximately 27 kDa, the expected molecular weight of sfGFP (**Fig. 8C, Supplementary Fig. 6**). LC-MS analysis of intact sfGFP-3TAG purified from cells expressing the PylRS/tRNA^Pyl^ pair from *Methanomethylophilus alvi* and *Methanohalophilus levihalophilus,* confirmed the introduction of Boc-Lys (**2**) or HO-Boc-Lys (**3**) into sfGFP-3TAG (**Fig. 8D**).The activity of the PylRS/tRNA^Pyl^ pair from *Methanohalophilus levihalophilus* is sufficient to support robust genetic code expansion with ⍺-amino acid and α-hydroxy acid Pyl analogs in *E. coli*.

A surprising result was the greater effectiveness of some PylRS/tRNA^Pyl^ pairs for incorporation of ⍺-hydroxy acid HO-Boc-Lys (**3**) compared to the ⍺-amino acid Boc-Lys (**2**). In particular, the PylRS/tRNA^Pyl^ pairs from Methanomicrobia JDFR-19 and *Methanogranum* 1D supported sfGFP expression only in the presence of HO-Boc-Lys (**3**). This finding is important because, unlike Boc-Lys (**2**), HO-Boc-Lys (**3**) generates an ester linkage within the protein backbone. Site-specific ester linkages can be exploited to post-translationally edit the protein backbone to introduce backbone elements that cannot otherwise be introduced, such as ꞵ^2^-amino acid linkages^18^ and reactive 1,3-diketones^19^. LC-MS analysis of intact sfGFP-3TAG purified from large-scale *E. coli* growths expressing PylRS/tRNA^Pyl^ pairs from JDFR 19 and *Methanogranum* 1 as well as *Methanohalobium* bin 14 and supplemented with ⍺-hydroxy acid HO-Boc-Lys (**3**) showed unambiguous evidence for the ester linkage (**Fig. 8C, Supplementary Fig. 7**). The selectivity for an ⍺-hydroxy acid substrate by the PylRS/tRNA^Pyl^ pairs identified here avoids well documented problems^16,54^ associated with metabolic conversion of ⍺-hydroxy acids into ⍺-amino acids that would circumvent post-translational backbone editing^18,19^.

## Clues to the pathway to genome-wide TAG reassignment

Based on the pattern of distribution and the functions of TAG codons in archaea that do, and do not incorporate Pyl, we suggest that the adoption of Genetic Code 34 may follow several steps (**Supplementary Fig. 8**).

First, the Pyl system may be acquired in some archaea via lateral transfer of a gene cassette and is associated primarily or exclusively with methylamine methyltransferases. Since the PYLIS is not required for Pyl incorporation, the Pyl machinery may incorporate Pyl at normal stop positions in these genomes. There may be an unknown means by which TAG is interpreted independently for each gene, but there is no evidence for this process at present. It is possible that there is transcriptional regulation of the Pyl machinery, potentially linked to methylamine availability. Supporting this are observations that a non-recoded Pyl archaean^55^ and a bacterium^24^ turn off the Pyl system when the organism is grown on a substrate other than trimethylamine, but the regulatory mechanism was not determined. The ability to turn off the Pyl machinery would eliminate unintended TAG readthrough, except when methylamines are available.

It has been suggested that Pyl incorporation at stop positions would not be highly detrimental and protein extension would be limited by backup stop codons (and non-functional proteins would be degraded)^11^. However, release factor 1 (RF1; recognizes TAG and TAA) is present in all complete and near-complete archaeal genomes examined and may compete with the Pyl tRNA for the TAG codon, reducing the instances of Pyl incorporation at stop positions. This situation is analogous to the ’ambiguous intermediate’ evolutionary model for recoding proposed for Candidate phyla radiation (CPR) bacteria and eukaryotes^7,56,57^.

The subset of genomes with many Category 4 and one or very few Category 1 genes (here designated as **<u>Group A</u>**) likely recently acquired the Pyl machinery and thus may represent the first stage of code transition. Interestingly all Group A genomes have ≥ 15% TAG genes, suggesting that high TAG stop codon use may be a barrier to code transition. Notably, the genomes that comprise Group A are phylogenetically very diverse, and include members of multiple phyla.

Over time, TAG codons would be introduced by mutation, leading to incorporation of Pyl within proteins. Simultaneously, potentially deleterious cases of TAG as a sense codon would decline due to mutation of the prior stop. Pyrrolysine is bulky, chemically reactive, and the ring nitrogen may introduce a positive charge into proteins at physiological pH, potentially making its incorporation unfavorable^42^. Thus, most mutations that introduce Pyl into organisms with Pyl machinery would be selected against. A subset of genomes have a small number of Category 1 TAG codons, but internal TAG codons occur both in methylamine methyltransferases and other genes. Genomes in this category typically have 5-15% TAG codons, possibly due to pressure to decrease use of TAG as a stop codon when the Pyl machinery is available. We infer that these genomes are in transition to Code 34, and designate genomes of this type as **<u>Group B</u>**. Code transition is deemed complete when TAG is never a stop codon. Transition may be inhibited by Category 4 codons, so the genomes most able to undergo full transition to Code 34 would be those with overall lower frequencies of TAG stop codons.

We designate as **<u>Group C</u>** those genomes with a majority of Category 1 genes, and few to no Category 4 genes, and classify them as having fully transitioned to Genetic Code 34. The Group C genomes typically have low TAG frequencies (≤ 5%), yet the JDFR-19 genome has 13% TAG and up to four Pyl within a single protein. Given that it takes time for the proteome to explore the potential value of Pyl incorporation and identify proteins in which Pyl is either neutral or beneficial, we suspect that JDFR-19 underwent code transition long ago. Notably, Pyl is orders of magnitude less abundant in any proteome than the next rarest amino acid (**Table S13**), probably because its incorporation is often detrimental to protein function.

Among Code 34 archaea, the only proteins that consistently contain Pyl are the methylamine methyltransferases. We surmise that, at some time in the past, dependence on anaerobic methylamine metabolism necessitated the availability of the Pyl system. Once present, this availability led to recoding the TAG codon as Pyl. The transition occurred only in methanogens, possibly because they have a narrow range of energetic substrates which can make them more dependent on Pyl than other more versatile microorganisms. Methanogens that adopted Code 34 are primarily obligate methylotrophic or methyl-reducing methanogens, thus most can only use methanol, methylamine and sometimes methyl-sulfides as sources of energy. The decreased utilization of TAG as a stop codon and its emergence as a Pyl codon in various essential genes likely occurs in methanogens living in conditions where methylamine is regularly the main source of energy (e.g. in saline environments where methylamines are produced from osmoprotectant degradation, or in the gut where methylamine precursors are abundant in certain diets). This transition could occur by two complementary processes. The constant requirement for pyrrolysine in methylamine methyltransferases may result in a loss of the Pyl machinery regulation. Independent of this loss, the constant occurrence of pyrrolysyl-tRNA synthetase and Pyl tRNA may lead to insertion of TAG in different essential genes. The insertion of TAG in essential genes would require that the Pyl machinery become constitutively expressed, even if the cells grow on compounds other than methylamines. Presence of TAG in essential genes and constitutive expression of the Pyl machinery would establish Code 34. This proposed mechanism of emergence of Code 34 in methanogens may also explain why it is not observed in bacteria, as they are generally more versatile and never methanogens. Thus, this is an instance of a metabolic process driving the adoption of a new genetic code.

We anticipate that TAG re-coding is a one way street, especially for archaea that have fully adopted Code 34. Loss of the Pyl machinery would result in loss of not only the ability to use methylamines, but also of the function of many Pyl proteins. As recoding was identified in individual genera, and only one archaeal family contains exclusively recoded genomes (**Fig. 4**), and given likely barriers to code reversal, the sporadic distribution suggests multiple independent code transitions. Thus, we infer that transition to Code 34 is facile. Interestingly, this parallels inferences of ease of code switching in some phages^6^ and contrasts with the monophyletic pattern of bacterial recoding^2,7^.

In summary, we find that the Archaea, previously not known to use more than a single genetic code, contains clades in which code switch has occurred such that the TAG stop codon, known to be fully reassigned in some eukaryotes and bacteria, is interpreted as pyrrolysine. We propose an evolutionary path to code switch and provide evidence that this code transition is relatively facile, occurring in many unrelated lineages. Finally, we demonstrate the biotechnological value of the newly identified pyrrolysine incorporation machinery.

## Methods

### Identification and annotation of Pyl- encoding Archaeal genomes

Genomes encoding Pyl were identified based on the presence of cellular machinery required for production and incorporation of this amino acid. We searched NCBI BlastP against the non-redundant protein database^58^ on December 1, 2023 with the pyrrolysyl-tRNA synthetase (PylRS) sequence from *Methanosarcina barkeri* and selected archaeal genomes with >40% amino acid sequence identity to the *M. barkeri* PylS sequence. Additionally, all genomes from each genus predicted to encode Pyl were downloaded based on taxon number from the NCBI website at https://www.ncbi.nlm.nih.gov/datasets/genome/?taxon= on December 1, 2023 (**Table S1**). For these genomes, Prodiga^l^^59^ v.2.6.3 was used to predict genes using default parameters (single genome mode, translation table 11), and hmmer^60,61^ v.3.3.1 was used via hmmscan against the NCBI-fam database^62^ v.13. The HMM gene annotation used to identify the Pyl pathway is as follows: TIGR03912 (PylSn), TIGR02367 (PylSc), TIGR03910 (PylB), TIGR03909 (PylC) and TIGR0391 (PylD). Genomes identified as encoding Pyl contain at least two of the Pyl-specific HMMs. Genomes with the following criteria were excluded from analysis: estimated completeness <60% and/or contamination (redundancy) >10%. CheckM^63^ v.1.1.6 was used for completion and contamination (redundancy) estimates. A recent Methanoglobus genome encoding Pyl was reported^28^ and released on February 25, 2024. This genome was added to the compilation of Pyl-encoding archaea and included for genomic analyses.

### Taxonomic distribution and phylogeny of Pyl machinery in Archaea

The reference species tree of Archaea was built from a concatenation of the sequences of 36 universally conserved proteins from the phylosift dataset^64^ together with sequences of L30 and S4 ribosomal proteins and of the RNA polymerase subunit A and B. Proteins were retrieved from the selected proteomes with HMM search^60,61^ (hmmer v3.3.2 [http://hmmer.org/]). They were aligned with MAFFT (v7.453) with the accuracy-oriented methods, L-INS-i^65^, trimmed with BMGE (v2) with the Blosum30 parameter^66^, and concatenated. A maximum likelihood tree was built with IQ-TREE (v2.0.6)^67^ using the LG+F+R10 model. This tree was used as a guide tree to run the Posterior Mean-Site Frequency (PMSF) model under the LG + C60 + F + G4 model implemented in IQ-TREE^68^ using a 1,000-replicate, ultra-fast bootstrap. Genome taxonomy was determined using GTDBTk 2.1.1^69^. Taxonomy, TAG categories and TAG% were mapped on the species tree using the ITOL server^70^.

PylBCDS proteins from archaea and bacteria were separately aligned using MAFF^65^ and trimmed with BMGE^66^ using the above mentionned parameters, and then concatenated. A maximum likelihood tree was built with IQ-TREE (v2.0.6)^67^ using the LG+R9. Presence/absence of PylSn and its fusion were mapped on the three, together with taxonomy using ITOL server^70^. Horizontal gene transfers were predicted by comparing PylBCDS topology with the reference species tree topology.

### Quantification of ORF TAG frequency

To quantify the mean TAG frequency of archaeal taxa, Pyl-encoding genomes were dereplicated for each phylum with dRep^71^ v.3.4.1 with default parameters. Then, Prodiga^l^^59^ v2.6.3 was used to predict ORFs using default parameters and the -d option used for nucleotide output. The Python script “find_tag_end.py” was used to retrieve all genes with an in-frame TAG codon. The TAG frequency was calculated as the total number of predicted ORFs containing an in-frame TAG codon, divided by the total number of predicted ORFs.

### Identification and quantification of genes by TAG codon category

To quantify TAG codon categories, we first extracted genes with in-frame TAG codons. NCBI genome coding sequence (CDS) files were used for gene identification. Prokka ^72^ V.1.14.5 was used for gene identification for genomes without NCBI CDS files. Each TAG codon was treated as a sense codon, and sequences containing TAG were extended until the nearest in-frame TAA or TGA stop codon. The resulting genes were then translated and annotated using the UniProt- SwissProt database version 2023_05 and the Uniprot Archaeal and Bacterial Reference Proteome databases^73^ version 2023_05 (November 8, 2023 release). TAG-containing genes were then grouped into one of the four designated categories, based on the location of the TAG codon: (1) Internal (2) Short extension (3) Long extension / short overlap / fusion and (4) Large out-of-frame overlap. Each gene was assigned to only a single category, using the lowest category number for which criteria were satisfied. Genes that were truncated due to being located at the end of a contig were excluded when extension length could not be determined. Category 1 (Internal) genes contain an internal TAG codon, and encode proteins for which the amino acid sequence before and after the TAG codon has ≥30% sequence identity with sequences from the UniProt databases. Additionally, for Category 1, the gene must also encode a protein that is >15 amino acids longer than if TAG was a stop. For Category 2 (short extension), readthrough of TAG results in a protein sequence extension of ≤15 aa.

Sequences overlapping an adjacent gene by ≥1 sense codon were excluded from Category 2. The Category 3 designation was assigned for genes if TAG readthrough resulted in (a) gene overlap of ≤60 bp with an adjacent gene, and / or (b) genes encoding a protein with >15 aa extension and/or (c) gene fusion (in-frame overlap) with an adjacent gene. Finally, TAG readthrough in Category 4 genes results in overlap of an adjacent gene by >60 bp, including overlap of an entire gene in a different reading frame or oriented in the opposite direction.

### Prediction of Pyl- containing proteins

We initially used the genetic code prediction tool, Codetta^48^, to identify Pyl proteins. By default, Codetta does not detect the recoding of TAG to Pyl. First, Codetta completes a six-frame translation of a genome nucleotide sequence that is aligned to profile HMMs. The software then recognizes recoding events when a codon has a specific nonstandard interpretation based on profile HMM alignments. For Codetta to complete this task, there must be a primary meaning that can be deduced from the alignments, for example, Codetta detected reassignment of AGG (a canonical arginine codon) to methionine^74^. When TAG is recoded to Pyl, TAG occurs in place of different codons such that there is no consensus for which codon it replaces. Thus, Codetta does not determine a single amino acid (re)interpretation for TAG, so it does not detect recoding. Identification of known Pyl proteins may be enabled with Codetta, but Pyl detection in this case is limited exclusively to methylamine methyltransferases because they are the only proteins with Pyl- containing profile HMMs that are available for Codetta. However, the alignment output produced by Codetta includes gene matches (profile HMMs) for each codon, including TAG. Using this Codetta alignment output, we identified Pyl-containing proteins when TAG is internal in a gene in a Code 34 genome. We implemented the alignment step using the Codetta v.2.0 codetta_align.py command with a compatible profile HMM database prepared with the hmmer^60,61^ v.3.1 hmmbuild command (--enone option) for the PGAP^62^ v.15 database, which includes TIGRFAM^75^ v.15.0 entries.

A secondary annotation for ORFs was performed by modifying the protein prediction software, Prodigal, to include TAG as a sense codon in the archaeal genetic code. The modified version of Prodigal is available at https://github.com/VeronikaKivenson/Prodigal and can be invoked using the -g 34 (translation table 34) option. Geneious was used to visualize and overlay the standard genetic code annotation and the TAG readthrough annotation. Sequences with in-frame TAG codon(s) were searched against Blast to identify protein matches. Searches for PYLIS structures were performed with Inferna^l^^76^ v.1.1.4 using covariance model(s) via cmsearch with the RFAM^77^ v.14.10 database. Prediction and annotation of Pyl- containing proteins was performed using the Bridges-2 computing resource^78^.

### Structural prediction of Pyl- containing proteins

Structures for homologous proteins were obtained from RCSB PDB^79^. Structure 4AY8 was used for MtaA^80^ and structure 7UTN was used for IscB^81^. The Visual Molecular Dynamics software package^82–84^ was used for visualization.

### Anaerobic cultivation

*Methanomethylophilus alvi* (formerly *M. alvus*) strain Mx-05^T^ (JCM 31474^T^) was grown as previously described^85^, with methanol and ruminal fluid. *Methanococcoides burtonii* strain ACE- M^T^ (DSM 6242^T^) was obtained from Deutsche Sammlung für Mikroorganismen und Zellkulturen (DSMZ) and grown on DSMZ Methanococcoides medium 141c (with 5 mM trimethylamine hydrochloride), with the following modifications: instead of Wolin’s mineral solution and Wolin’s vitamin solution, ATCC trace mineral supplement (MD-TMS) and ATCC Vitamin Supplement (MD-VS) were used. *M. burtonii* was grown at room temperature in 15 mL Hungate tubes with a headspace of 85% N_2_, 10% CO_2_, 5% H_2_ gas in Anaerobe Systems anaerobic chamber (AS- 150).

### Sample preparation for LC-MS/MS

20 mg cell pellets were resuspended in 100 mM ammonium bicarbonate, pH 8.0 and subjected to mechanical lysis by bead beating with 0.15 mm zirconium oxide beads for five min (Geno/Grinder 2010; SPEX). Resulting cell lysates were adjusted to 4% sodium dodecyl sulfate (SDS)/10 mM dithiothreitol (DTT), incubated at 90°C for 10 min to denature and reduce proteins, and pre-cleared by centrifugation at 21,000 x g. Samples were adjusted to 30 mM iodoacetamide (Sigma) and incubated in the dark at room temperature for 20 min to alkylate cysteine residues. The alkylated proteins were further cleaned and prepared for proteolytic digestion via the protein aggregation capture (PAC) method^86^. In brief, 300 μg of hydrophobic magnetic beads (1 micron, SpeedBead Magnetic Carboxylate; GE Healthcare UK) and acetonitrile (ACN) were added to protein samples for a final concentration of 80% ACN to induce protein aggregation on the beads. Samples were incubated for 20 min at room temperature. The aggregated proteins were placed on a magnet and the supernatant was removed. The pelleted beads were subjected to a wash with 1 mL of ACN, followed by 1 mL of 70% ethanol. The final wash buffer was removed from tubes and the beads were resuspended in 200 µL of 100 mM ammonium bicarbonate for in-solution proteolytic digestion. Protein amounts were quantified by corrected absorbance (Scopes) at 205 nm (NanoDrop OneC; Thermo Fisher). After protein quantitation, samples were digested using either MS-grade GluC Endoproteinase or trypsin (Thermo Scientific) with a 1:75 protease:protein (wt:wt) ratio at 37°C, shaking overnight at 600 rpm. An additional round of protease was added for a second 3-h digestion period at 37°C, shaking at 600 rpm. The resulting tryptic peptides were filtered on 10 kDa MWCO filter plate (AcroPrep Advance, Omega 10 K MWCO) at 1,500 × g and adjusted to 1% formic acid before quantification by NanoDrop OneC.

#### LC-MS/MS

3 µg of peptide solution were analyzed by automated 1D LC-MS/MS analysis using a Vanquish ultra-HPLC (uHPLC) system plumbed directly in-line with a QExactive- Plus mass spectrometer (Thermo Scientific) with an in-house-pulled 100 µm inner diameter nanospray emitter (packed to 15 cm with 1.7 µm Kinetex C18 reverse-phase resin (Phenomenex)). Peptides were loaded, desalted, and separated by uHPLC under the following conditions: sample injection followed by 100% solvent A (98% H_2_O, 2% acetonitrile, 0.1% formic acid) from 0 to 30 min to load and desalt, a linear gradient from 0 to 30% solvent B (70% acetonitrile, 30% water, 0.1% formic acid) from 30 to 220 min for separation, a column wash from 0 to 100% solvent B from 220 to 255 min, and 100% solvent A from 255 to 275 min for column re-equilibration. Eluting peptides were analyzed with the following MS settings for global searches: positive mode, data-dependent acquisition, top-20 method; mass range 400–1500 m/z; MS and MS/MS resolution 70 and 17.5 K, respectively; MS/MS loop count 20; isolation window 1.8 m/z; isolation offset 0.3 m/z; charge state exclusion of unassigned, +1, +6–8 charges.

For targeted searches, parallel reaction monitoring inclusions lists were constructed to detect peptides of interest in the predicted pyrrolysine containing proteins. An in-silico digest of the proteins was conducted with UniPept^87^ using either GluC or trypsin as the protease with a minimum peptide length of six amino acids. Based on the predicted peptides from the *in silico* digest, peptides were selected for the PRM measurements if they were classified as any of the following peptide types: (1) the first proteolytic peptide in the protein, (2) the proteolytic peptide containing the pyrrolysine residue, (3) the first proteolytic peptide downstream of the pyrrolysine-containing peptide, and (4) the last proteolytic peptide in the sequence. For the peptides of interest, the PRM inclusion list included the monoisotopic precursor m/z of the peptide including a static modification of carbamidomethylation (+57.021) of cysteine and a charge state (z) of +2. No variable modifications of oxidation (+15.995) of methionine, deamidation of (+0.984) of asparagine and glutamine, or methyl loss (-14.016) for pyrrolysine were considered for the inclusion list. A maximum of 50 entries were monitored for each PRM measurement. The same LC-MS/MS method was used as the global measurements.

### Proteomics data analysis

Global database searches were conducted with Proteome Discoverer 3.0 (Comet/Percolator)^88,89^ against custom organism specific protein databases constructed from all predicted proteins in the respective, as well as common mass spectrometry contaminants.

Protein sequences were included with the predicted pyrrolysine residue represented with “X” as well as the sequence version that terminated in a stop codon. For all database searches, the peptide mass tolerance was set at 20 ppm. Accepted modifications included static modifications of carbamidomethylation (+57.021) of cysteine residues and pyrrolysine (+237.148) of “X” residues. Database searches were performed with variable modifications of oxidation (+15.995) of methionine, deamidation of (+0.984) of asparagine and glutamine residues, and a methyl loss (-14.016) for pyrrolysine (X) residues, and a maximum of three variable modifications allowed per peptide. Semi-proteolytic digestion of peptides was allowed for either trypsin or GluC depending on the sample, and a 1% peptide-level FDR threshold was applied.

Preliminary confirmation of pyrrolysine expression was based on (1) a unique peptide match in the global datasets that passed the 1% peptide-level FDR threshold for the pyrrolysine containing proteins or (2) a detected precursor mass in the PRM measurement that was within 10ppm of the monoisotopic precursor m/z of the peptides of interest. Peptides were considered unique if the amino acid sequence was not predicted in any other protein for the respective organism. To more confidently confirm pyrrolysine expression, MS/MS spectra from all peptides of pyrrolysine-containing proteins were manually inspected and validated. A MS/MS spectrum of a peptide was considered for evidence of pyrrolysine expression if each of following criteria was met: (1) Precursor ion (MS1) mass error <10 ppm, (2) Fragmentation ion (MS/MS) mass error <0.02 Da and (3) Location of the fragment ion along the peptide sequence, with either direct sequencing of the fragment ending in pyrrolysine or fragment ions that were longer than the pyrrolysine-containing fragment.

In addition to the database searches, we searched the data using a de novo-assisted database search workflow using PEAKS StudioX^90^ to recognize any of the standard amino acids at the TAG=pyrrolysine/stop position (represented as “X” in the database). This workflow will recognize the 20 standard amino acids if they were detected at the Pyl position in the MS/MS spectrum. The same modifications as previous database searches were included and the parent and fragment ion mass error tolerances were set to ± 10 ppm and ±0.02 Da, respectively. De novo only parameters were left at default settings with average local confidence (ALC) scores of >80% and de novo sequence tags displayed if at least six amino acids were shared with the database sequence.

#### Construction of plasmids encoding PylRS/tRNA^Pyl^ pairs

Plasmids encoding the PylRS/tRNA^Pyl^ pairs evaluated in Fig. 8 were derivatives of the previously reported pMEGA vector (parent backbone Addgene Plasmid #200225). pMEGA plasmids containing each PylRS homolog (**Table S14**) were ordered as circular, dsDNA plasmids from Twist Bioscience. Each one was then modified to eliminate the M. alvi pylT gene and replace it with the one corresponding to each PylRS homolog (**Table S14**). Briefly, each circular pMEGA vector obtained from Twist was linearized using oligonucleotides P1 and P2 (**Table S14**). PCR using the Q5® High-Fidelity 2X Master Mix (NEB M0492S) was performed for 30 cycles according to manufacturer protocols. The PCR reaction was subject to a PCR clean up with the QIAquick PCR Purification Kit (Qiagen, catalog # 28104). The concentration of purified, linearized products was determined by absorbance at 260 nm using a NanoDrop ND- 1000 Spectrophotometer. Next, 33.3 ng of each purified, linearized pMEGA plasmid was combined with 100 ng of a gBlock containing the appropriate pylT sequence (**Table S14**) in a 20 μL Gibson Assembly reaction using HiFi DNA Assembly Master Mix (NEB, catalog #E2621L). The reaction mixture was incubated at 50 °C for 1 h to generate a family of circular pMEGA vectors containing the coding sequence for the appropriate pylT.

The circularized pMega plasmids from the previous step were then transformed into NEB 5- alpha competent E. coli (NEB, catalog # C2987H) as follows. Frozen stocks of cells were thawed on ice for 10 min. Upon thawing, the entirety of the previous Gibson Assembly reaction was added to cells and incubated on ice for 30 min. Cells incubated with plasmid were then subjected to heat shock at 42 °C for 30 sec and placed on ice for 2 min. 350 μL of Super Optimal broth with Catabolite repression (S.O.C.) outgrowth medium (NEB, catalog # B9020S) was added to cells and cells were incubated at 37 °C for 1 h with shaking at 220 rpm. Agar plates containing spectinomycin were inoculated with 50 μL of transformed cells and grown overnight at 37°C. 3 single colonies per construct were picked and inoculated into liquid cultures containing 5 mL LB + spectinomycin and grown for 16 h at 37°C. Pure plasmid was isolated from 5 mL cultures using Qiaprep Spin Miniprep Kit (Qiagen, catalog # 27106) and sequences were confirmed by whole plasmid sequencing with Primordium Labs. The new pMEGA plasmids were double transformed with a pET22b-3TAG-GFP into BL21(DE3) cells following the transformation protocol detailed below.

### Transformation protocol

BL21(DE3) E. coli (NEB, catalog # C2987H and C2527H, respectively) were transformed in accordance with manufacturer protocols with some modifications as follows. Frozen stocks were thawed on ice. Upon thawing, 100 ng of each relevant plasmid was added. After a 30 min incubation on ice, cells were heat-shocked for 30 sec at 42 °C and allowed to recover on ice for 2 min. Following, 350 μL of Super Optimal broth with Catabolite repression (S.O.C.) was immediately added and cells were allowed to recover at 37°C with shaking for 1 h before plating 50 µL on LB agar plates containing the appropriate antibiotics. Plates were incubated overnight at 37 °C.

### Expression assays of sfGFP-3TAG containing Boc-Lys or HO-Boc-Lys

Starter E. coli cultures were grown overnight in 5 mL of LB Miller (AmericanBio, Catalog #AB01201) in 15 mL culture tubes supplemented with carbomycin and spectinomycin at 37°C. The next morning, 500 µL of the saturated overnight culture was added to 50 mL LB in a 250 mL baffled flask with carbomycin and spectinomycin and grown at 37 °C to an OD_600_ of 0.6.

Prior to cultures reaching OD600 0.6, the appropriate amount of each monomer (stored as 25 mM stocks) were allotted into the wells of a black, clear bottom, 96-well plate (1 mM final concentration). Once cultures reached an OD_600_ of 0.6 (roughly 3 h), protein expression was induced by addition of 1 mM IPTG. The culture was transferred into the wells of the plate to a total volume of 200 µL in each well. A Breathe-Easy® sealing membrane (Sigma Z380059) was placed over the 96-well plate and the plate was loaded into a BioTek Synergy HTX microplate reader with no lid. OD_600_ and F_528_ values (ƛ_ex_ = 485 nm) were measured every 10 min for 20 h.

The plate was maintained at 37°C and was shaken during this time. Points represented in **Fig. 8B** represent three biological replicates where three random colonies were picked on a plate from a single transformation.

### Expression and Purification sfGFP-3TAG containing Boc-Lys or HO-Boc-Lys

Starter cultures of 5 mL of Miller’s LB Broth (AmericanBio, catalog # AB01201) supplemented with carbomycin and spectinomycin were inoculated with a single colony of BL21(DE3) E. coli cells harboring pET22b-sfGFP-3TAG-6xHis alongside pMEGA-JDFR19-PylRS-pylT, pMEGA- JDFR19-PylRSCTD-pylT, pMEGA-B74G9-PylRSCTD-pylT, pMEGA-LMO1-PylRS-pylT, pMEGA-Bin14-PylRS-PylRS-pylT, pMEGA-1D-PylRS-pylT, pMEGA-Levihalophilus-PylRS-pylT, or pMEGA-MaPylRS and grown overnight for 16 h at 37°C with shaking at 220 rpm until the culture was saturated. The starter culture was used to inoculate a 100 mL expression culture in a 1:100 dilution of Miller’s LB Broth supplemented with 1 mM Boc-Lys (Combi-Blocks, CAS # 2418-95-3) or 1 mM HO-Boc-Lys (Accela, CAS # 111223-31-5) with carbomycin and spectinomycin. The expression culture was grown at 37°C with shaking at 220 rpm to an OD_600_ of 0.6 at which point it was induced with 1 mM IPTG and grown for 16 h under the same conditions. The expression culture was harvested by centrifugation at 4,300 x g at 4 °C for 30 min. The resulting cell pellet was suspended in 10 mL of Lysis Buffer (50 mM sodium phosphate pH 6.8 and 300 mM NaCl) containing half a tablet of cOmplete, mini EDTA-free ULTRA protease inhibitor cocktail (Sigma-Aldrich, St. Louis, MO). The cell suspension was disrupted by sonication on ice (Branson Sonifier 250, 5 cycles of 30 sec pulse at 50% duty cycle and microtip limit of 5 followed by 30 sec pause). The cell lysate was cleared by centrifugation at 10,000xg at 4 °C for 20 min. TALON® Metal Affinity Resin (1 mL) (Takara Biosciences, catalog # 635504) was equilibrated with Lysis Buffer, added to the cleared cell lysate, and incubated on a rotisserie at 4 °C for 1 h. The TALON® resin-lysate mixture was then passed through a gravity flow Poly-Prep Chromatography Column (Bio-Rad Laboratories, Hercules, CA). Non-specifically bound proteins were removed by washing the TALON® resin with 10 mL of Lysis Buffer. The 6xHis- tagged protein was eluted by washing the TALON® resin with 2 mL of Elution Buffer (50 mM sodium phosphate pH 6.8 and 250 mM imidazole). The purified protein was quantified using absorbance at 280 nm, snap frozen as single-use aliquots, and stored at -80 °C. Protein yields are reported in **Table S15**.

### Intact Protein LC-MS for GCE

LC-MS analysis of all protein samples were performed on an Agilent 1290 Infinity II HPLC connected to an Agilent 6530B QTOF AJS-ESI. The mobile phase for LC-MS was water and acetonitrile with 0.1% (v/v) formic acid at a flow rate of 0.4 mL/min. Each protein sample was injected onto an Poroshell 300SB-C8 column (2.1 x 75 mm, 5 µM, room temp, Agilent) and separated using a linear gradient from 5% acetonitrile for 0 to 2 min and ramping to 95% acetonitrile over 7.5 min, and then washing with 95% acetonitrile for 2 min. The following parameters were used during acquisition: Fragmentor voltage 225 V, gas temperature 300 °C, drying gas flow 10 L/min, sheath gas temperature 350 °C, sheath gas flow 11 L/min, nebulizer pressure 35 psi, skimmer voltage 65 V, Vcap 5000 V, 1 spectra/s.

## Data Availability and Code Availability

Proteomic data from this study have been deposited into the ProteomeXchange repository with accession numbers: ProteomeXchange-PXD053523; MassIVE-MSV000095193. Reviewers can use the FTP download link with the MassIVE accession number to access the files. We will provide login information to reviewers upon submission and release this data publicly upon publication. The modified version of Prodigal with Genetic Code 34 that has TAG reassigned to Pyl is available at https://github.com/VeronikaKivenson/Prodigal and the python script (find_tag_end.py) is available at https://github.com/VeronikaKivenson/pyrrolysine.

## Acknowledgments

We thank Jean-François Brugere, Johnathan Rylee, and Petar Penev for helpful discussions. This work was supported by a Tory Burch Foundation Fellowship to VK at the Innovative Genomics Institute. This work used Bridges-2 at the Pittsburgh Supercomputing Center through allocation BIO230230 to VK, from the Advanced Cyberinfrastructure Coordination Ecosystem: Services & Support (ACCESS) program, which is supported by National Science Foundation grants #2138259, #2138286, #2138307, #2137603, and #2138296. LTR was supported by the NSF Graduate Research Fellowship Program (DGE-1752814). Experimental validation of PylRS activity for ⍺-amino and ⍺-hydroxy acids was supported by the NSF Center for Genetically Encoded Materials (C-GEM; CHE 2002182). Funding was provided by the Chan-Zuckerberg Initiative Foundation, Technology Enabled Biological Carbon Capture and Sequestration (CZIF2022-007203) and National Science Foundation, Collaborative Research: TRTech-PGR Track (2334028) to JFB. SG wishes to thank the Miller Institute at UC Berkeley for a Visiting Professorship. GB was supported by Agence Nationale de la Recherche (ANR-19-CE02-0005). Proteome work done at ORNL was supported by a U.S. Department of Energy Genome Sciences Program project (ERKPA50).

## Author contributions

V.K. and J.F.B. conceived and designed the study. V.K. performed codon analysis, identification of Pyl proteins, and anaerobic cultivation of *M. burtonii*. Experimental validation of PylRS activity for ⍺-amino and ⍺-hydroxy acids was performed by N.X.H. and L.T.R. with oversight by A.S. Proteomics experiments and data analysis was conducted by S.L.P with oversight by R.L.H. Phylogenetic analysis and discussion was conducted by G.B. and S.G. Structural prediction of Pyl-containing proteins and bioinformatics software development was performed by A.K. Anaerobic cultivation of *M. alvus* was performed by K.F. with oversight by G.B. The manuscript was written by V.K. and J.F.B. with input from all authors. All authors read and approved the manuscript.

## Ethics declarations

### Competing interests

The Regents of the University of California have a patent pending related to this work on which V.K., L.T.R., N.X.H., A.S. and J.F.B. are inventors.

## Supplementary Figures

**Fig. S1.**
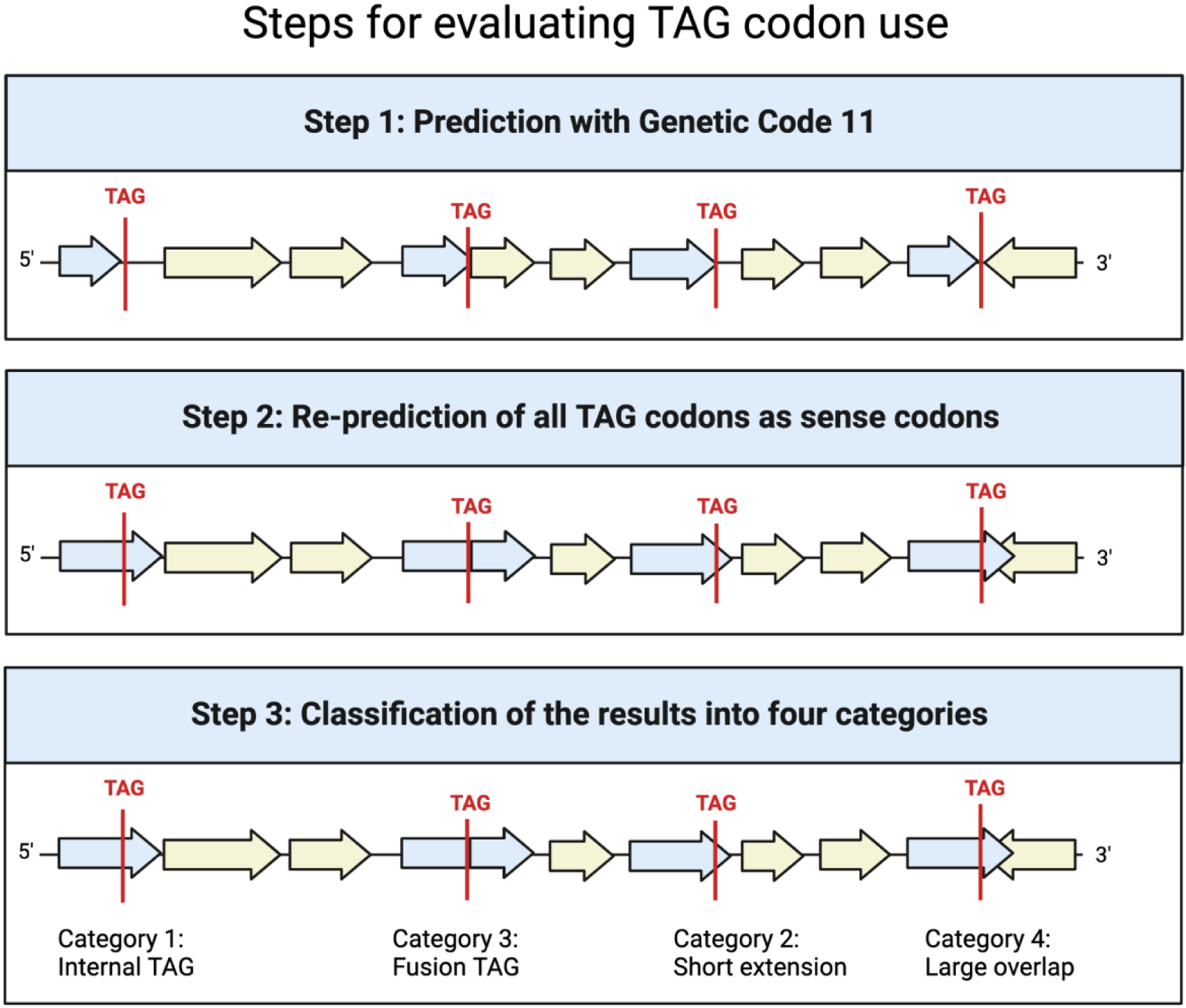
Steps for assigning TAG codon into different categories. Genes in blue contain a TAG codon, while other genes are shown in yellow.

**Fig. S2.**
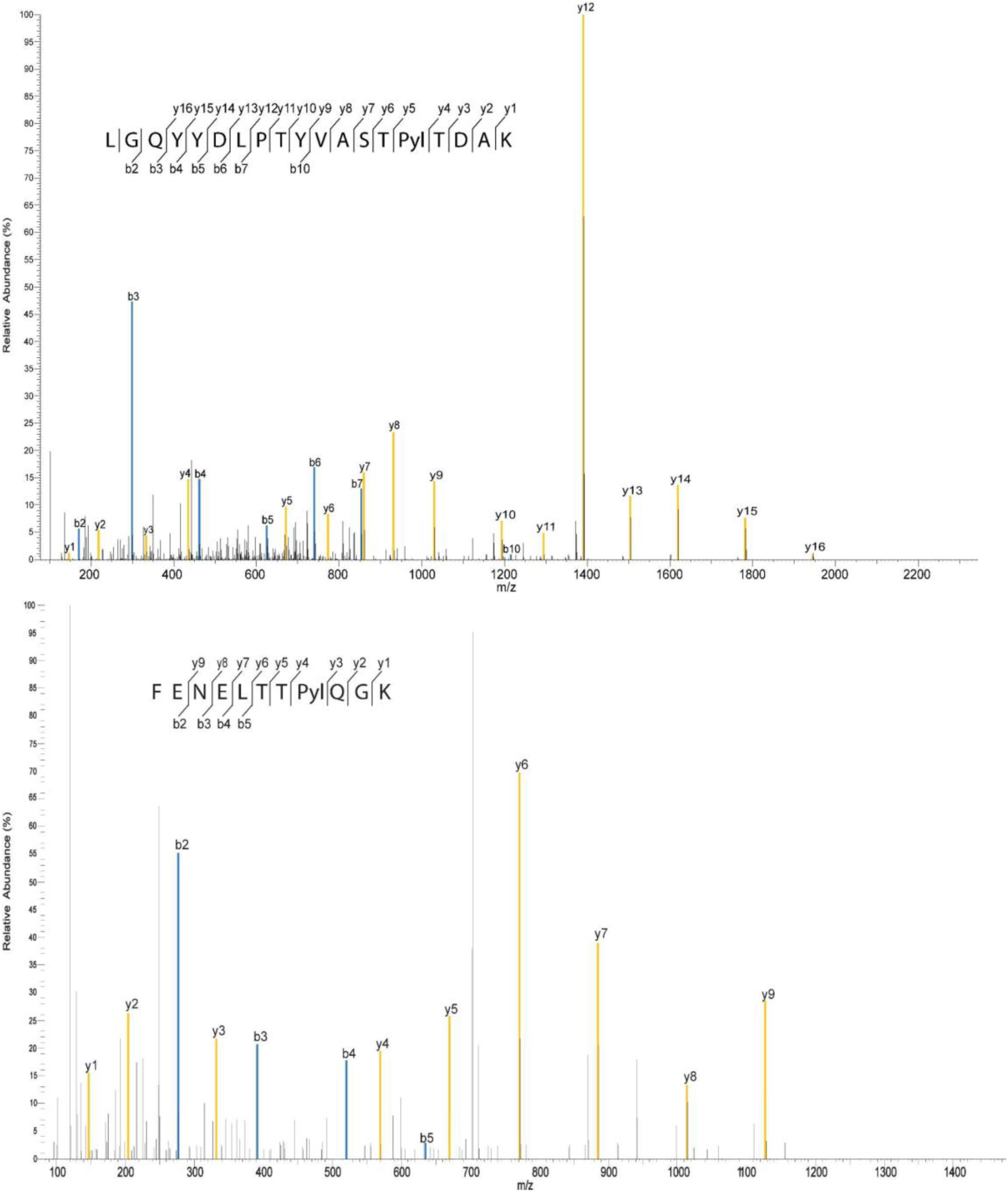
MS/MS spectrum of two Pyl-containing proteins. The blue peaks represent observed b-ions and the yellow-peaks represent observed y-ions. Black peaks were not identified as b- or as y-ions. Observed b- and y-ions are indicated in the peptide sequence ladder as well as in the MS/MS spectrum **a,** (Upper panel) MS/MS spectrum of Pyl-containing tryptic peptide of *M. burtonii* deacetylase from the global DDA measurements. **b,** (Lower panel) MS/MS spectrum of Pyl-containing tryptic peptide for *M. burtonii* trimethylamine methyltransferase 2 from the global DDA measurements.

**Fig. S3.**
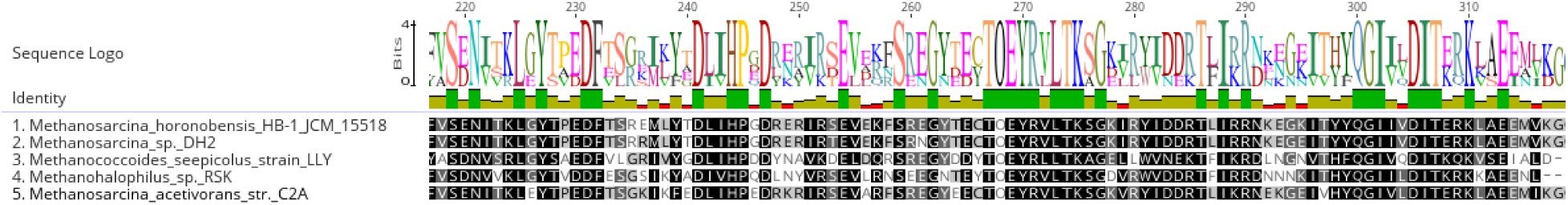
Alignment of protein (GAF domain-containing / PAS family protein); Pyl (O) is at position 269.

**Fig. S4.**
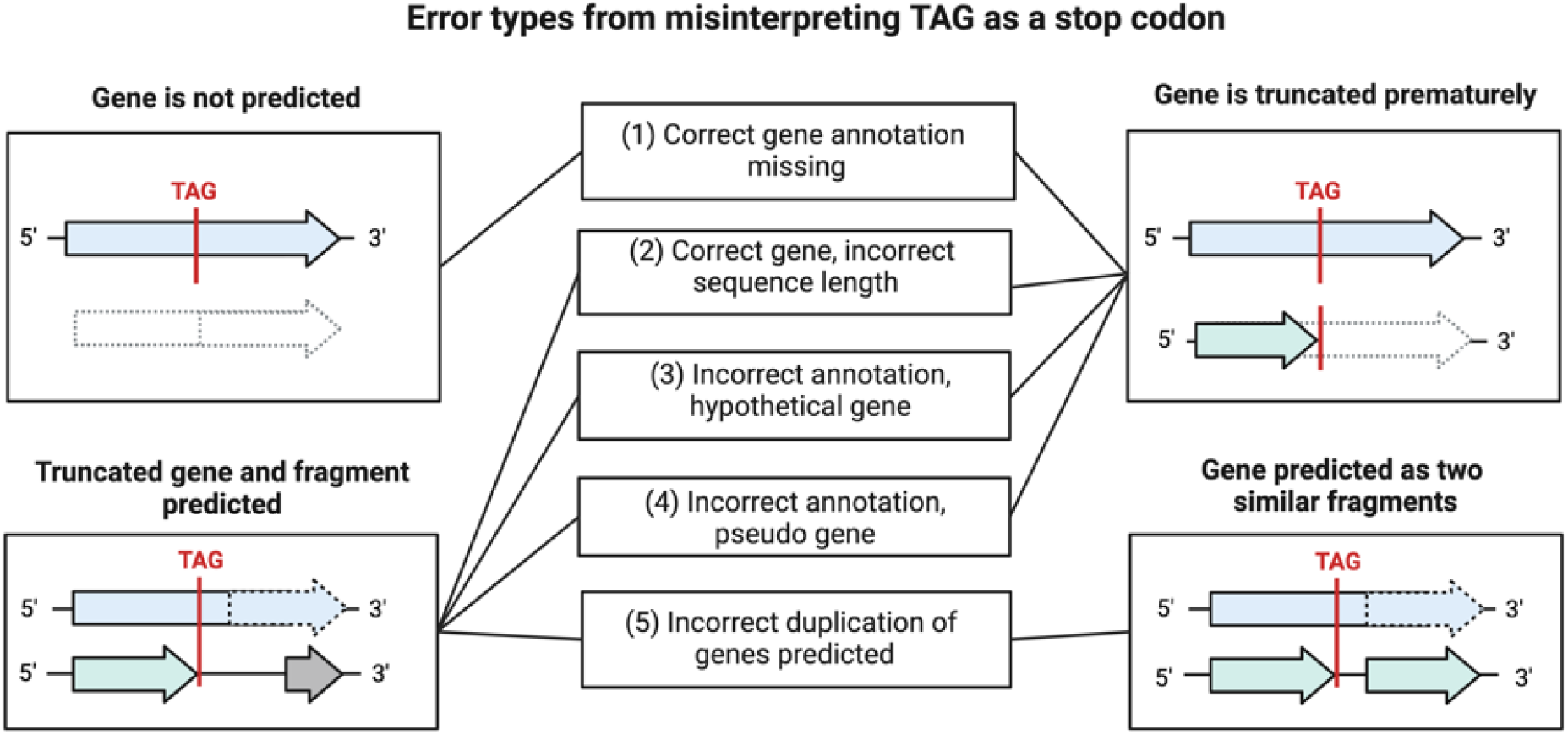
Different types of misannotation for Pyl proteins resulting from misinterpretation of TAG as a stop codon.

**Fig. S5.**
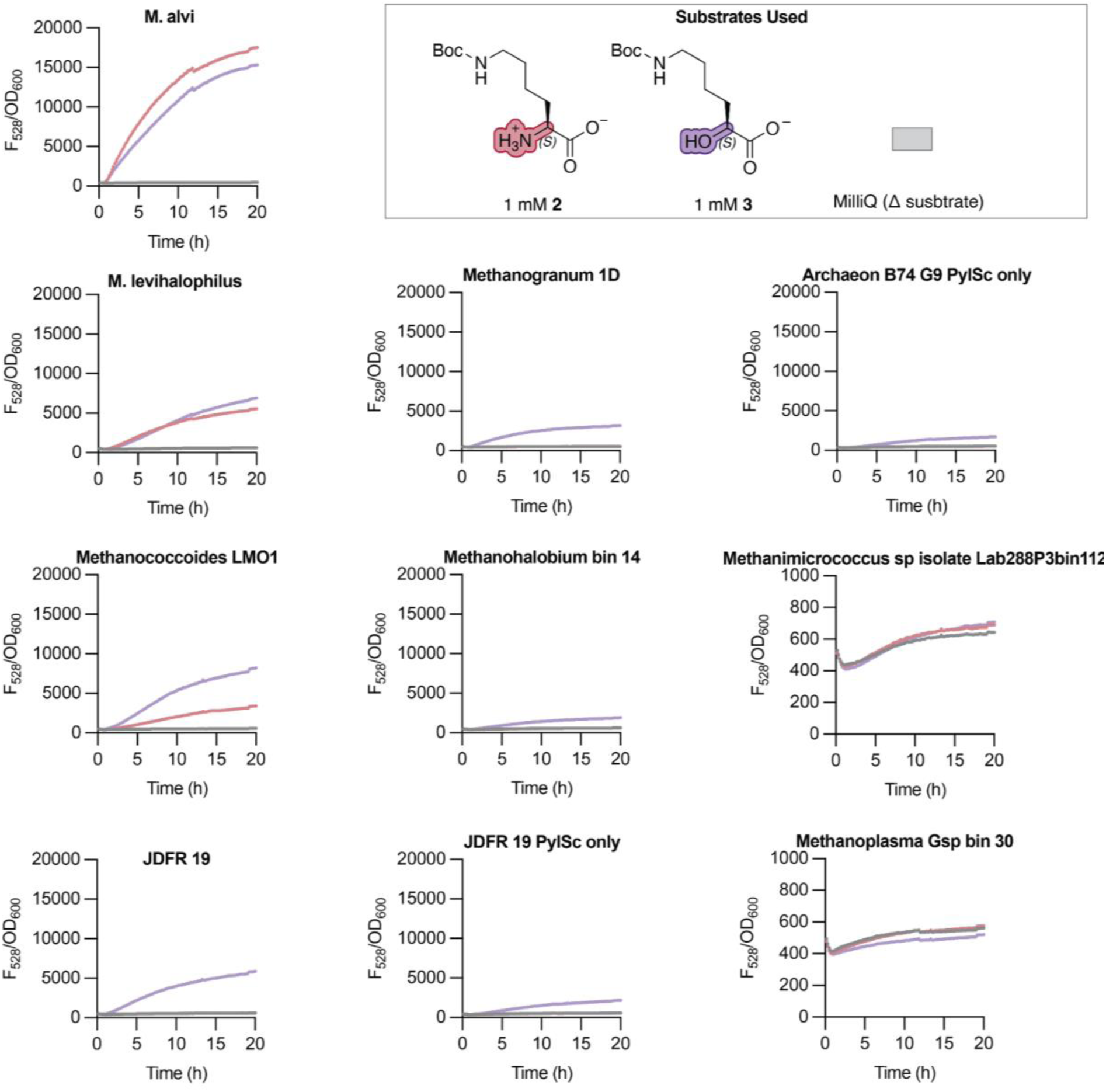
Time-dependent changes in 528 nm emission (F_528_) and cell density (OD_600_) of BL21(DE3) *E. coli* harboring a pMega plasmid expressing the indicated PylRS/tRNA^Pyl^ pair and a reporter plasmid encoding sfGFP-3TAG and grown in the presence of 1 mM BocK (monomer 1) or ⍺-HO-BocK (monomer 3). Growth and expression conditions are described further in Supplementary Information.

**Fig. S6.**
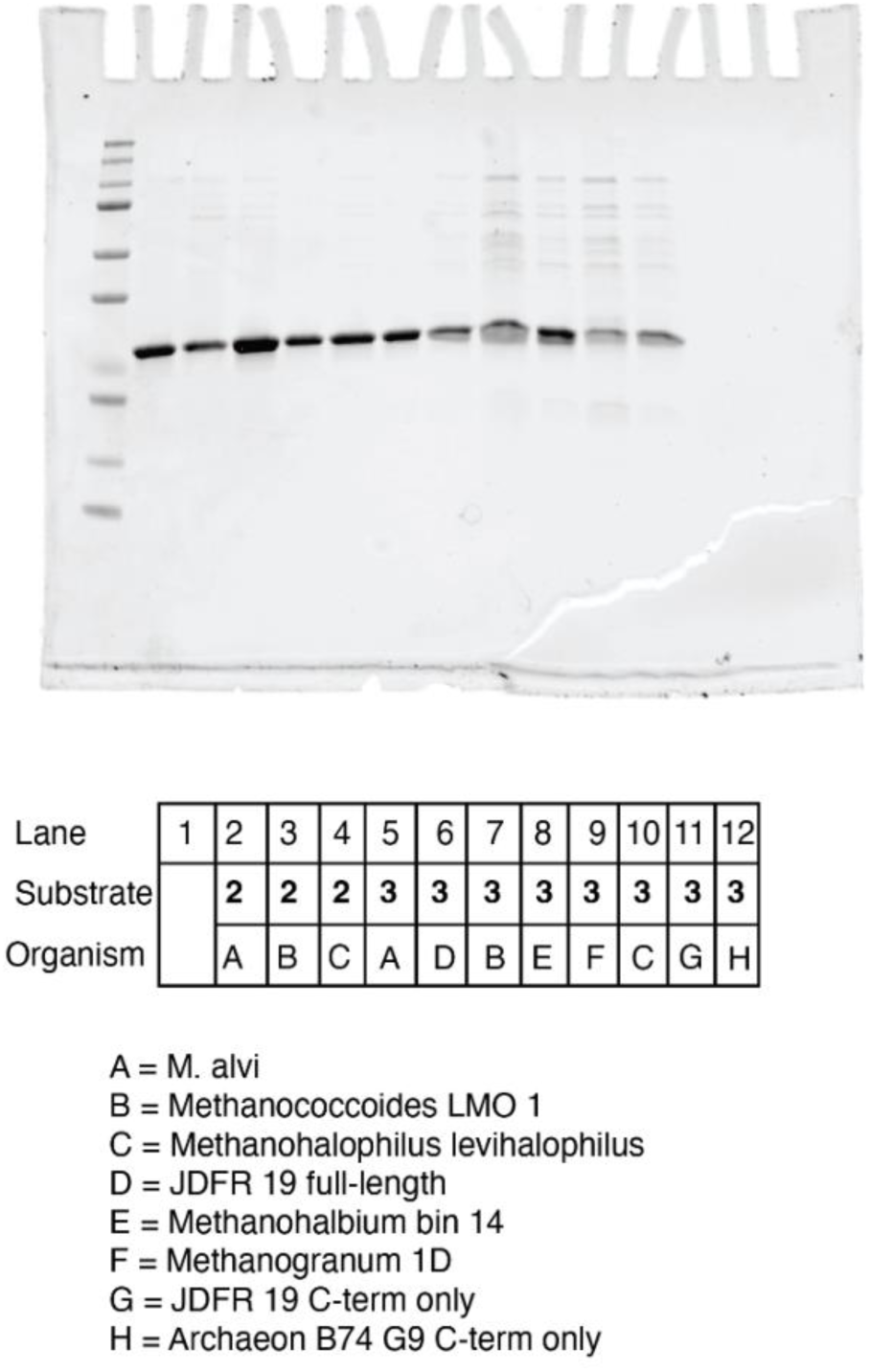
Uncropped SDS-PAGE gel of lysates from BL21(DE3) cells transformed with a pMega plasmid encoding the indicated PylRS/tRNA^Pyl^ pair and a reporter plasmid encoding sfGFP-3TAG.

**Fig. S7.**
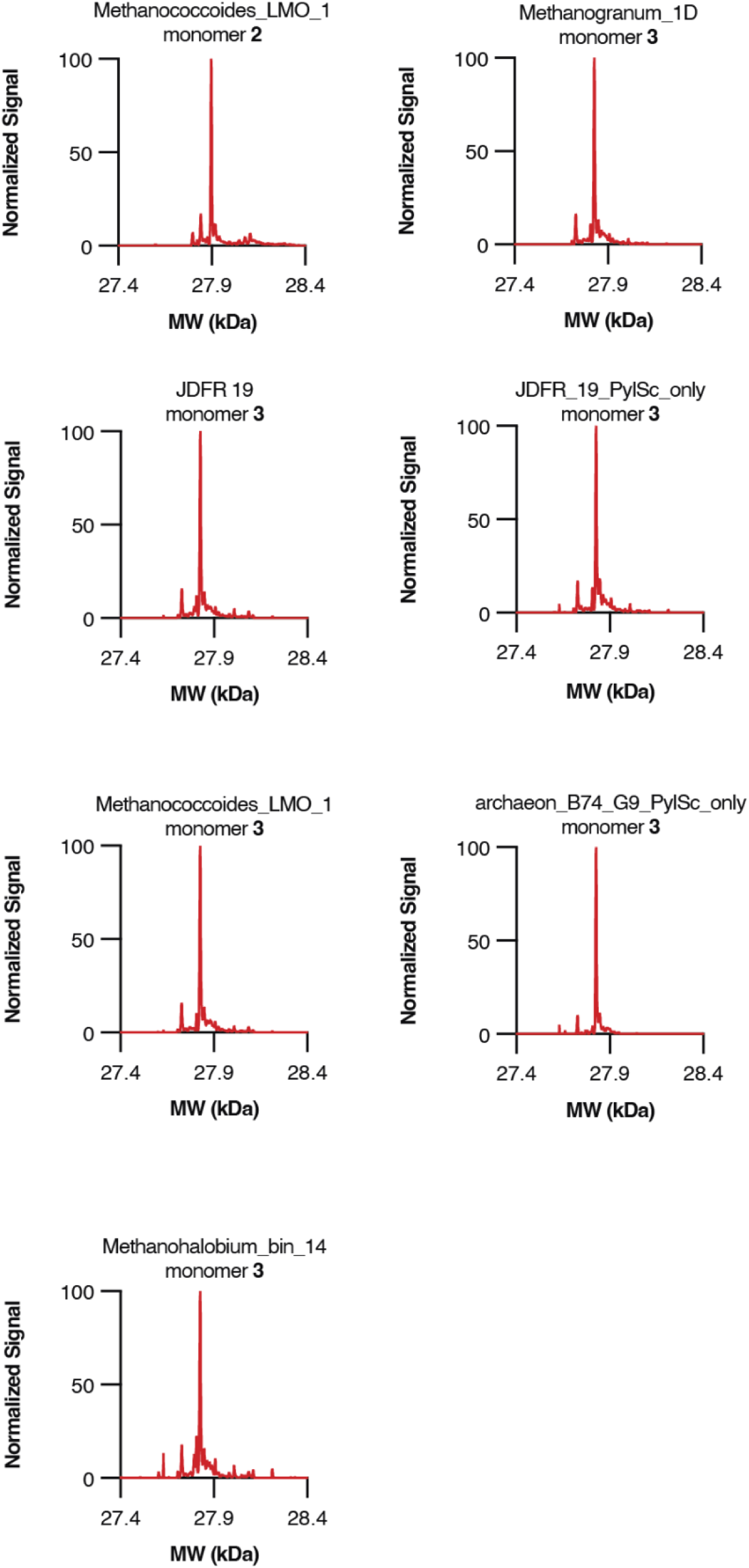
LC-MS characterization of sfGFP-3TAG produced in BL21(DE3) cells supplemented with either Boc-Lys (monomer 2) or ⍺-HO-Boc-Lys (monomer 3). Shown are the deconvoluted mass spectra of purified sfGFP-3TAG isolated from BL21(DE3) *E. coli* transformed with a pMega plasmid encoding the indicated PylRS/tRNA^Pyl^ pair and a reporter plasmid encoding sfGFP-3TAG.

**Figure S8.**
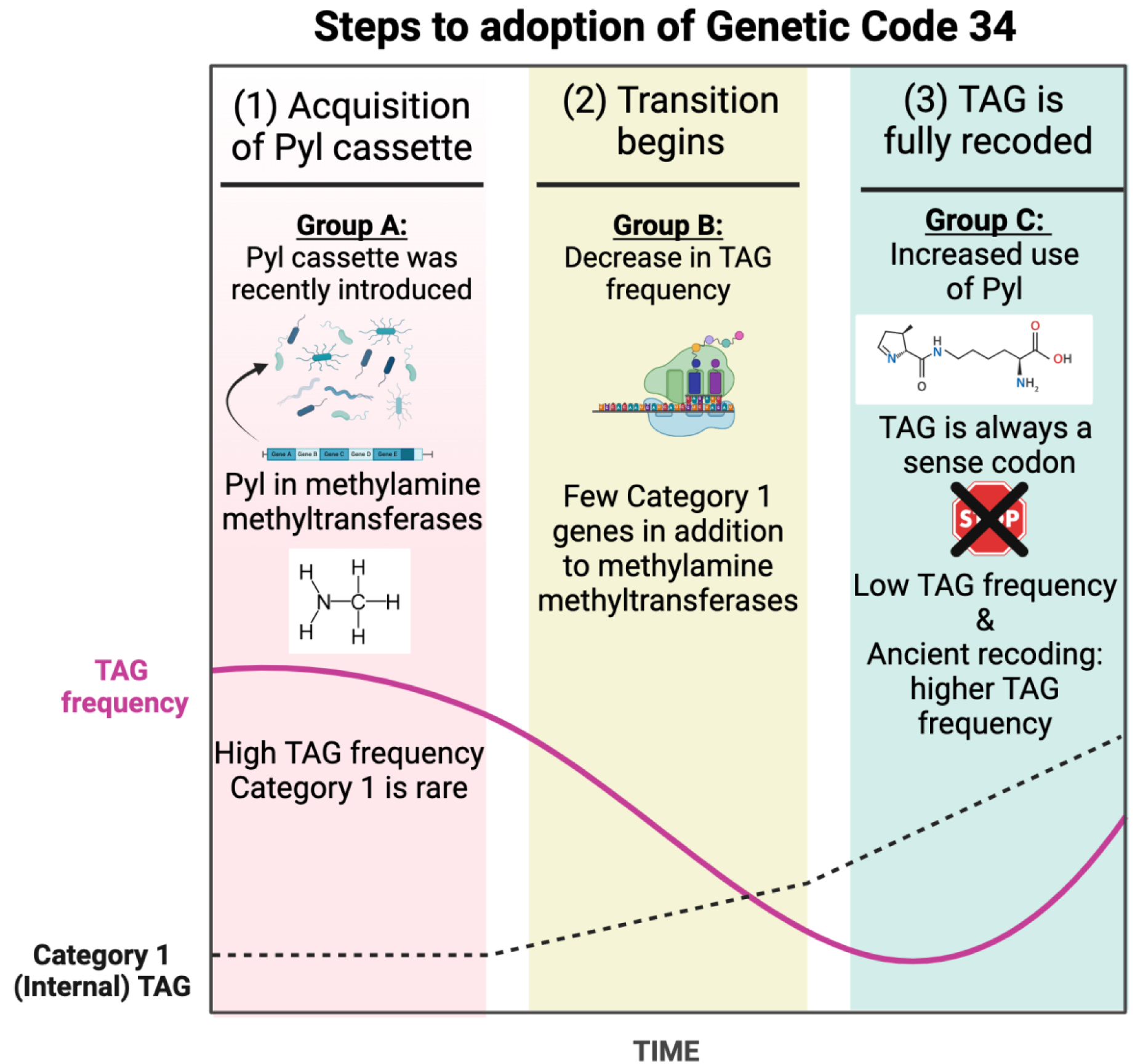
A schematic representation of the proposed mechanism for the evolution of genome-wide TAG recoding. Organisms that have recently aquired the Pyl cassette still retain their prior genomic distribution of TAG, with frequent use of TAG as a stop codon and little to no incorporation of TAG into genes (column 1). The presence of the Pyl cassette exerts a strong evolutionary pressure against the use of TAG as a stop codon, leading to a large decrease in TAG frequency. At the same time, TAG coding for Pyl begins to appear inside genes, potentially through neutral mutation (column 2). Finally, organisms well-adapted to the Pyl cassette no longer use TAG as a stop codon but their TAG frequency increases despite this due to the accumulation of TAG coding for Pyl inside genes (column 3).

## SUPPLEMENTARY TABLES

https://docs.google.com/spreadsheets/d/19mk-sypDhi80z_soNEorhwmjfLQ8-3rZEDywxOZUgb8/edit?usp=sharing

**Table_S1:** List of previously reported archaeal groups reported to encode Pyl and corresponding references.

**Table_S2:** List of Pyl genomes including their accession numbers and environment or source.

**Table_S3:** Taxonomic summary of Pyl taxa and their taxon number, and genomes encoding Pyl machinery by taxonomic group.

**Table_S4:** Search parameters for genes encoding Pyl-containing proteins examined to identify PYLIS.

**Table_S5**: Quantification of genes with TAG codons by designated codon categories.

**Table_S6:** Compilation of the results from both the global and targeted measurements of expressed proteins from *M. alvi*, showing coverage of Pyl-containing proteins.

**Table_S7:** Compilation of the results from both the global and targeted measurements of expressed proteins from *M. burtonii*, showing coverage of Pyl-containing proteins.

**Table_S8:** Summary of the new proteins found to contain Pyl and confirmed by proteomics for M. alvi and M. burtonii.

**Table_S9:** List of putative Pyl-containing proteins predicted from genomes with Code 34.

**Table_S10:** Putative Pyl-containing proteins predicted via TAG codon alignment within HMM models for conserved proteins using Codetta.

**Table_S11:** List of putative Pyl proteins from the JDFR-19 genome.

**Table_S12:** List and sequences of CRISPR-Cas, IscB, TnpB, and casposon proteins with Pyl residues.

**Table_S13:** Codon usage quantification for the JDFR-19 genome.

**Table_S14:** Gene fragments, oligos, and plasmids used in *in vivo* GCE studies.

**Table_S15:** sfGFP yield from expression and purification sfGFP-3TAG containing Boc-Lys or HO-Boc-Lys.

